# Whole-genome Omics delineates the function of CCM1 within the CmPn networks

**DOI:** 10.1101/2023.07.11.548554

**Authors:** Jacob Croft, Brian Grajeda, Luis A Aguirre, Liyuan Gao, Johnathan Abou-Fadel, Victor Sheng, Jun Zhang

**Author notes:** All correspondence: Jun Zhang, Sc.D., Ph.D., Department of Biomedical Sciences, Texas Tech University Health Science Center, 5001 El Paso Drive, El Paso, TX 79905.

## Abstract

**Introduction:** Cerebral cavernous malformations (CCMs) are abnormal dilations of brain capillaries that increase the risk of hemorrhagic strokes. Mutations in the KRIT1, MGC4607, and PDCD10 genes cause CCMs, with mutations in CCM1 accounting for about 50% of familial cases. The disorder exhibits incomplete penetrance, meaning that individuals with CCM may appear normal initially, but once symptoms manifest, their brains have already suffered irreversible damage. Compromised blood-brain barrier (BBB) is crucial in regulating the flow of substances between the blood and the central nervous system, which can result in hemorrhagic CCMs. Progesterone and its derivatives have been studied for their impact on maintaining BBB integrity. CCM2 interacts with CCM1 and CCM3, forming the CCM signaling complex (CSC), which connects classic and non-classic progesterone signaling to establish the CmPn signaling network, vital in preserving BBB integrity.

**Methods:** The study aimed to explore the relationship between CCM1 and key pathways of the CmPn signaling network, utilizing a toolset comprising three mouse embryonic fibroblast lines (MEFs) with distinct CCM1 expression levels. Omics and systems biology analysis were performed to investigate Ccm1-mediated signaling within the CmPn signaling network.

**Results:** The findings suggest that CCM1 plays a critical role in controlling cellular processes in response to different progesterone-mediated actions within CmPn/CmP signaling networks, partly by regulating gene transcription. This function is crucial for preserving the integrity of microvessels, indicating that targeting CCM1 could hold promise as a therapeutic approach for this condition.

## Introduction

Cerebral Cavernous Malformations (CCMs) are a common type of brain vascular malformation that results from abnormally dilated capillaries in the brain. This condition leads to increased risk of hemorrhagic strokes. CCMs are an autosomal dominant disorder with three known genes, KRIT1 (CCM1), MGC4607 (CCM2), and PDCD10 (CCM3) as causes of familial CCMs (1). Mutations in the CCM1 gene account for about 50% of familial CCM cases (2, 3). One distinct characteristics of this genetic disorder is its incomplete penetrance, meaning that most people with the mutated gene are asymptomatic. However, when symptoms do occur, they often result in irreversible brain damage. To study the cellular functions of CCM1, researchers have established isogenic mouse embryonic fibroblasts (MEFs) from wild type (WT) and Ccm1-knockout (KO) mice and created Ccm1-knockin (KI) fibroblasts by stable infection of Ccm1-KO fibroblasts with a lentiviral vector encoding human KRIT1 (4–8).

The Blood-Brain Barrier (BBB) is a crucial interface between the blood and the central nervous system that regulates the flow of substances between the two (9, 10). The BBB integrity is maintained by two major apparatus, adherens junctions (AJs) and tight junctions (TJs), which are formed by different molecules but are functionally and structurally linked (11). Inflammatory events are a major cause of BBB disruption (12, 13), and steroids are often used therapeutically for their anti-inflammatory properties in human conditions, including BBB disorders (11, 14, 15), although the efficacy of steroids in this context is still a topic of debate (11). Similar to estrogen and glucocorticoids, the effects of progesterone (PRG) and its derivatives, progestins, on BBB integrity have been investigated, despite a lack of understanding of their off-target effects via their corresponding steroid receptors (16). PRG binds to nuclear receptors (nPR) in classic PRG actions and membrane progesterone receptors (mPRs) in non-classic PRG actions (17–23).

Recent findings indicated that CCM2, which anchors with both CCM1 and CCM3 to form the CCM signaling complex (CSC) (1, 24). The CSC, in turn, can couple both classic nuclear progesterone receptor (nPRs) and non-classic membrane receptor (mPRs) mediated progesterone (PRG) signaling to form the CSC-mPRs-PRG-nPRs (CmPn) signaling network in nPR(+) cells, and the CSC-mPRs-PRG (CmP) signaling network in nPR(-) cells (23, 25, 26). The CmPn/CmP signaling network was recently found to play a crucial role in maintaining BBB integrity through its impact on nPR(-) microvascular ECs in both in vitro and in vivo conditions, suggesting the essential roles of CmPn/CmP signaling networks in maintaining microvascular integrity (26, 27). In order to better understand the cellular roles of CmPn/CmP signaling networks during angiogenesis and tumorigenesis, our research team has been conducting multi-omics studies using different cellular and animal models to investigate the key role of CCM1 within the CmPn signaling network. Specifically, we have focused on the relationship between CCM1 and other key components of the CmPn signaling network (nPRs, mPRs) (23, 25). This study investigated the differentially expressed protein (DEP) profiles in mouse embryonic fibroblasts (MEFs) with varying levels of CCM1 protein expression in response to progesterone (PRG) actions. The DEPs were categorized into up-regulated and down-regulated genes in each CCM1 genotype under progesterone (PRG) actions. The non-biased, whole-genome scale analysis allowed us to quantitatively measure Omic dynamic changes in response to progesterone (PRG) actions and define Ccm1-mediated signaling within the CmPn signaling network. The findings from our study demonstrate the crucial functions of the CCM1 protein in the signaling networks of CmPn/CmP. They provide insights into the potential transcriptional regulatory pathways associated with this relationship and emphasize the therapeutic potential of targeting CCM1 for treating health conditions related to the CmPn/CmP networks. These results will greatly contribute to understanding the molecular mechanism of CmPn/CmP signaling networks, particularly the signaling mediated by CCM1 in these networks.

## Materials and Methods

### Cell Culture, treatment, sample preparations and data acquisition

#### Cell Culture and Treatment

All three MEF cell lines (CCM1-WT, CCM1-KO, CCM1-KI/96) were cultured at 37°C and 5% CO2 in Dulbecco’s modified Eagle’s medium (DMEM) supplemented with 10% fetal bovine serum (FBS), 2 mM glutamine, and 100U/mlpenicillin/streptomycin as described (28). In this work, we defined progesterone (PRG) action as the combination of progesterone and mifepristone (PRG+MIF). To compare the effects of progesterone (PRG) action, MEF cells at 80% confluence were treated with different conditions: vehicle control (ethanol/DMSO, VEH), progesterone (PRG) actions (PRG+MIF; 20µM each), or no treatment (Untreated). These treatments were applied to examine and compare the impact of progesterone (PRG) action, as previously described (25, 26).

#### Cell collection, protein sample preparations and data generation

After the cells from various treatment groups were rinsed and harvested, proteins were extracted from cell lysis using a lysis buffer. Following the extraction of proteins, a purification process utilizing well-established methods was performed to ensure the acquisition of high-quality proteins for subsequent protein data acquisition (24–26, 29). To obtain Omic data, we employed established protocols (24) and utilized liquid chromatography tandem mass spectrometry (LC-MS/MS) for peptide analysis. Subsequently, either MaxQuant or Skyline was employed for data processing in the next phase.

#### Proteomic data acquisition and processing

Using the proteomics data we obtained, we conducted an analysis of the protein expression patterns in Mouse Embryonic Fibroblasts (MEFs) with three different genotypes of CCM1: Ccm1-WT, Ccm1-KO, and Ccm1-KI. Additionally, we examined how these protein expression patterns responded to progesterone treatments. The protein expression patterns of MEFs with three genotypes of CCM1 (Ccm1-WT, Ccm1-KO, and Ccm1-KI) and their response to progesterone (PRG) treatments were analyzed through the use of our acquired proteomics data. For the proteomic data analysis, Sequest was employed to search the UniProt reference protein database, with specific parameters defined, including a fragment ion mass tolerance of 0.020 Da and a parent ion tolerance of 10.0 PPM. In Sequest, we applied a fixed modification of carbamidomethyl on cysteine residues, along with variable modifications of oxidation on methionine residues and acetyl on the N-terminus. To ensure the reliability of peptide and protein identifications based on MS/MS data, Scaffold was utilized. Only identifications surpassing a probability threshold of 95.0% (for peptides) and 99.0% (for proteins), as determined by the Peptide Prophet and Protein Prophet algorithms respectively, were considered valid. The false discovery rate was determined by utilizing the decoy database. In cases where MS/MS analysis alone could not distinguish between proteins, they were grouped together following the principles of parsimony. MaxQuant software was utilized for protein quantification, and protein abundance was expressed as intensity-based absolute quantification (iBAQ) values. To ensure data quality, low-quality identifications and contaminants were filtered out from the resulting peptide and protein identifications. Differential protein expression analysis was carried out using DEP, Percolator, or ProteinQuant tools to identify proteins that were differentially expressed (DEPs) between the various CCM1 genotypes and their respective responses to the different conditions.

#### RNAseq data acquisition and processing

All RNA-seq data were produced using Illumina HiSeq 2000 platform; followed by RNAseq raw data process. The workflow of RNAseq data process involved the removal of rRNA reads from the raw data, followed by the filtration of low-quality reads (with more than 20% of base qualities below 10), and the removal of reads with adapter sequences and unknown bases (more than 5% N bases). Total clean reads for all samples were over 99.5%; 60-80% of reads were mapped to reference genomes. All clean reads were then assembled into unigenes, annotated for function, and expression levels and SNPs were calculated for each sample. Differential expressed genes (DEGs) were identified, and clustering analysis and functional annotations were performed to gain further insights. The detailed acquisition of RNAseq data was carried out as previously described (24). The objective was to uncover the similarities and differences among two genotypes (Ccm1-KO, and Ccm1-KI/96) under either mPR-specific treatments or vehicle control. The consistency in differential expression was analyzed using a Python comparison script. For the data analysis, the first step was to remove reads that were mapped to ribosomal RNAs to obtain the raw data. Next, we filtered low-quality reads (reads with more than 20% of bases with a quality score lower than 10), reads with adapter sequences, and reads with unknown bases (more than 5% N bases) to obtain the clean reads. We then assembled the clean reads into Unigenes, and annotated their functions. The expression levels and SNPs of each sample were then calculated. Finally, we identified DEGs (differentially expressed genes) among the samples and performed clustering analysis and functional annotations.

### Omics, Bioinformatics Analysis, and Systems Biology

#### Pathway enrichment analysis

To investigate the gene-gene interactions within the pathways related to the function and signaling of the CCM1 gene, we performed pathway enrichment analysis. During this analysis, proteins were grouped based on significant peptide evidence and annotated using Gene Ontology (GO) terms (30) and the Kyoto Encyclopedia of Genes and Genomes (KEGG) (31). GO and KEGG pathway analyses are performed to identify enriched pathways associated with differentially expressed genes. The pathway analysis data is integrated using methods like Protein-Protein Interaction Network (PPI) analysis, Gene Set Enrichment Analysis (GSEA), and comparative omics to identify relationships between proteins and aggregates of signaling pathways based on set intersections across multiple sets (32–34).

#### Systems biology analysis

The proteins are grouped based on significant peptide evidence and annotated with Gene Ontology (GO) terms (30) and Kyoto Encyclopedia of Genes and Genomes (KEGG) database (31). GO and KEGG pathway analyses are performed to identify enriched pathways associated with differentially expressed genes. The pathway analysis data is integrated using methods like Protein-Protein Interaction Network (PPI) analysis, Gene Set Enrichment Analysis (GSEA), and comparative omics to identify relationships between proteins and aggregates of signaling pathways based on set intersections across multiple sets (32–34).

The study conducted basal-level statistical analysis to determine the differential expressions of various components. T-tests were performed on differentially expressed proteins in multiple clusters from various sample comparisons. The study included three pairs for varying levels of CCM1 and three pairs for the effect of different genetic backgrounds under PRG actions. Each experimental set was biologically replicated three times. The t-tests were based on quantitative values without multiple test correction, a significance level of p < 0.05, fold change by category, a quantitative method, and normalization. The results were collated into an Excel sheet, which also included basal-level statistics for each pathway component.

#### ML-aided Transcriptional Factors (TF) prediction analysis

To enhance the efficiency of transcription factor prediction, we developed our approach to combine the utilization of Evolutionary Scale Modeling (ESM) with a cost-sensitive Support Vector Machine (SVM), with the following three key steps: **1**). ***Evolutionary Scale Modeling (ESM).*** ESM is an advanced transformer-based language model designed specifically for proteins. It has undergone training on an extensive corpus containing 250 million protein sequences (35). The main objective of this training was to equip the model with the ability to directly predict protein structure, function, and various properties from individual sequences using masked language modeling. Due to its inherent flexibility, ESM is exceptionally well-suited for fine-tuning a wide range of tasks that necessitate protein sequences as input. At the heart of our methodology lies the utilization of ESM to generate highly representative encodings of protein sequences, which are subsequently employed as input for our prediction model. In this study, we specifically utilized employ ESM-2, a cutting-edge protein language model that has demonstrated superior performance compared to all tested single-sequence protein language models in various structure prediction tasks. Moreover, ESM-2 enables the prediction of atomic resolution structures (36). Its capabilities in comprehending and predicting protein interactions and functions are unparalleled. **2**). ***The Cost-sensitive SVM***. SVM is a classifier known for its robustness in distinguishing classes within the feature space by identifying an optimal hyperplane. However, traditional SVMs can face challenges when dealing with imbalanced datasets, as they do not take into account the varying costs associated with misclassifying examples from minority classes. To tackle this issue, we adopt a cost-sensitive SVM approach, where the misclassification costs for each class are assigned based on their inverse proportion to their frequencies. This approach ensures a more equitable treatment of minority classes and addresses the limitations of traditional SVMs (37). 3). ***Experimental Setup and Data Preprocessing.*** Our ML-assisted experimental setup involves conducting experiments using a 10-fold cross-validation strategy on a dataset comprising 3738 protein sequences. The dataset consists of 2784 samples classified as Non-Transcription Factors (NTFs) and 954 samples classified as Transcription Factors (TFs). To ensure consistency across sequences, we apply truncation or padding techniques to achieve a standardized length of 1000 amino acid residues. This preprocessing step ensures that all sequences are of the same length for further analysis and model training.

## Results

### Differential expressed proteins between mouse embryonic fibroblasts (MEFs) with different CCM1 genotypes

The experiment analyzed differentially expressed protein profiles in MEFs with three distinct CCM1 genotypes, classified as low endogenously expressed CCM1 (CCM1-WT), total depletion of CCM1 (CCM1-KO), and ectopically overexpressed CCM1

(CCM1-KI/96). These genotypes exhibited significantly different expression levels of CCM1 protein.

#### Differentially expressed protein (DEP) profiles in the MEFs with different CCM1 genotypes

In this comparative proteomic analysis, a Venn diagram was used to identify differentially expressed proteins (DEPs) in three CCM1 genotype comparative pairs: wild type MEFs with low endogenously expressed CCM1 genotype (CCM1-WT) vs ectopically overexpressed CCM1 genotype (CCM1-96), CCM1-depletion genotype (CCM1-KO) vs CCM1-WT genotype, and CCM1-overexpressed (CCM1-96) genotype vs CCM1-WT genotype. The analysis revealed that 559 DEPs were identified in the first genotype-pair, with 205 proteins (19.4%) being unique to this comparison (Fig. 1A, blue circle). Additionally, 487 DEPs were identified in the second genotype-pair, with 120 DEPs (11.3%) being specific to this comparison (Fig. 1A, yellow circle). Furthermore, 621 DEPs were defined in the third genotype-pair, with 194 DEPs (18.3%) being exclusive to this comparison (Fig. 1A, pink circle). Out of the 1059 total identified proteins, 68 (6.4%) were shared by all three-comparison genotype-pairs, 113 (10.7%) were shared by the first (blue)/second (yellow) genotype-pairs, 173 (16.3%) were shared by the first (blue) /third (pink), and 186 (17.6%) were shared by the second (yellow)/third (pink) genotype-pairs (Fig. 1A). The DEPs were further categorized as up-regulated (red-color bar) and down-regulated (blue-color bar) in each comparative CCM1 genotype-pair. The results showed that more upregulated genes were exclusively identified in the first genotype-pair, while more downregulated genes were observed in the second and third genotype-pairs (Fig. 1B). A heatmap displayed the notable differences in DEPs among MEFs with the three different CCM1 genotype comparative pairs, indicating their significantly differential expression of protein (DEP) profiles (Fig. 1C).

**Figure 1.**
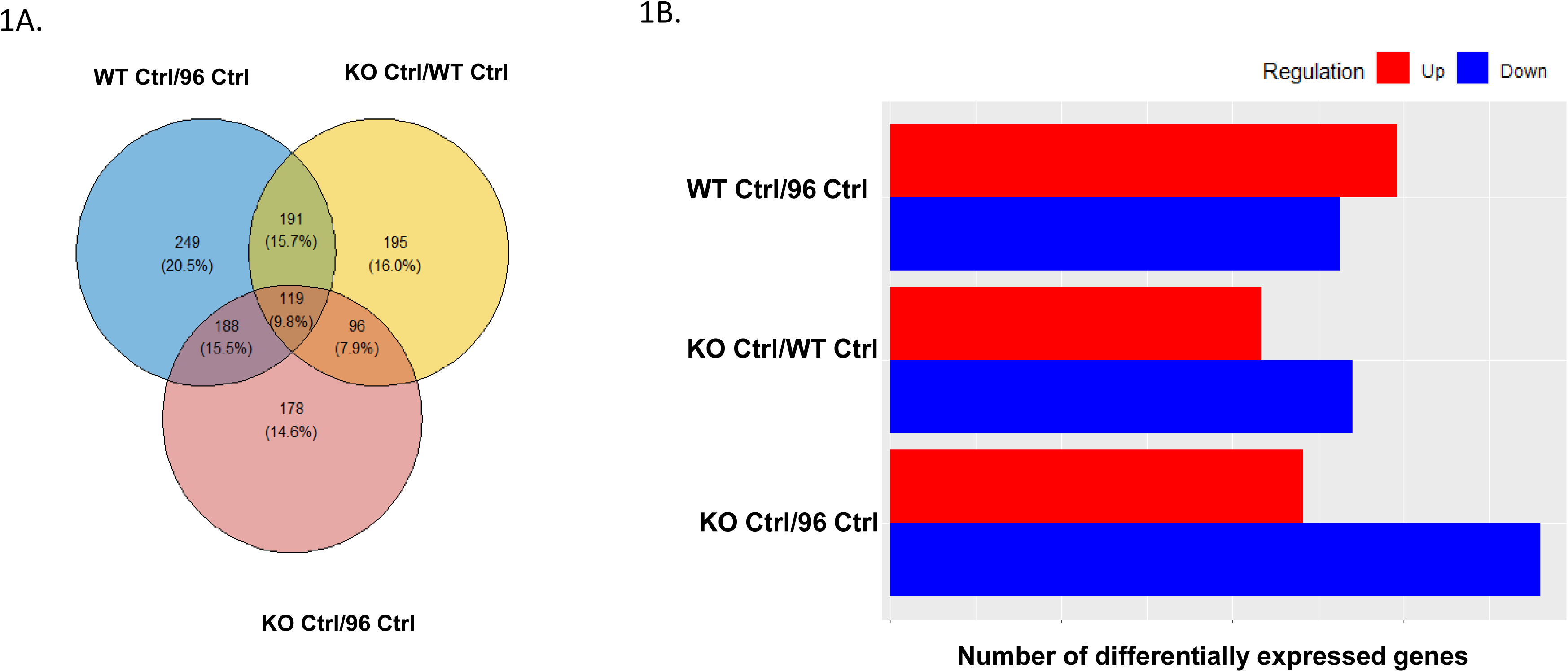
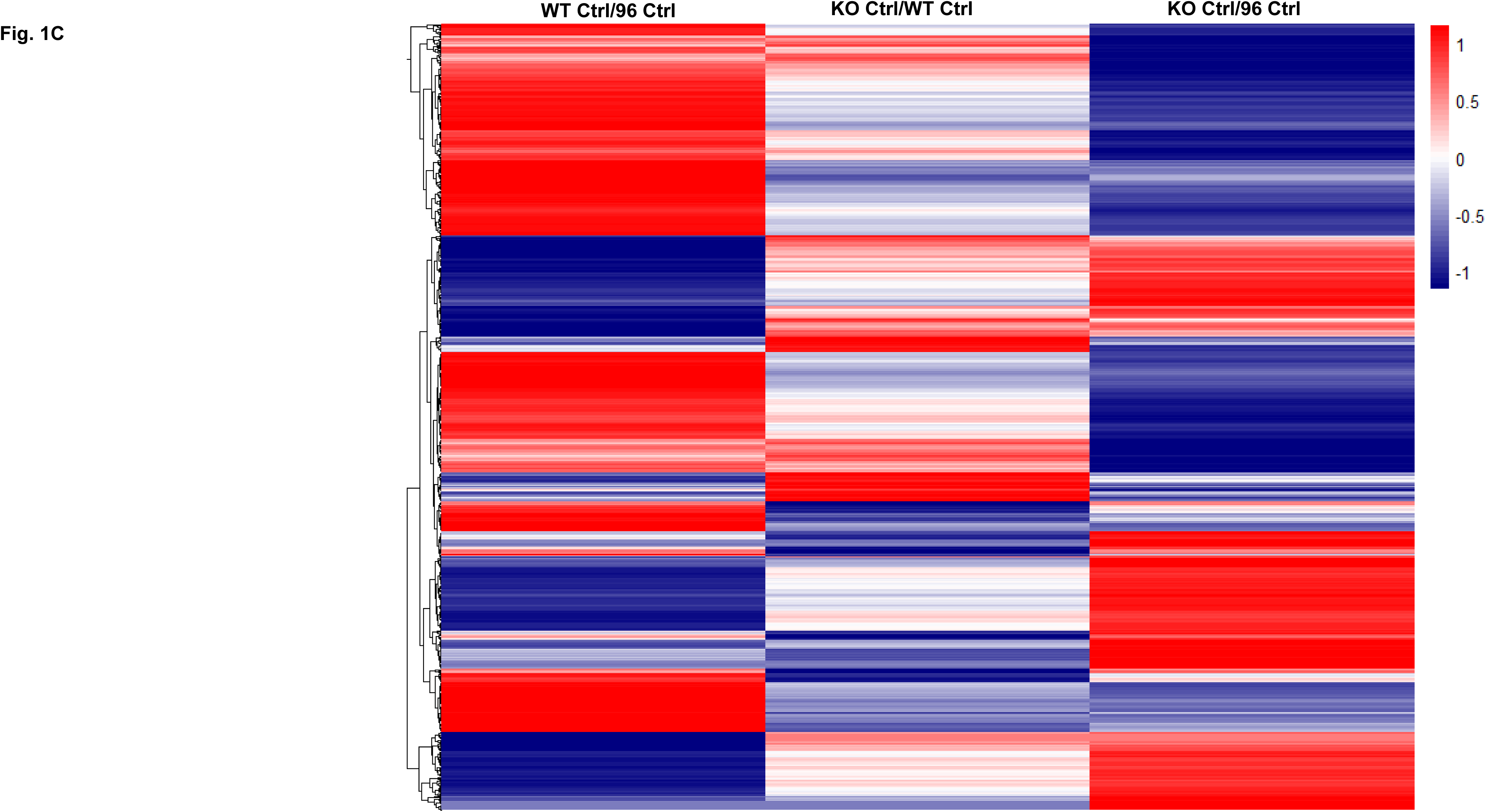
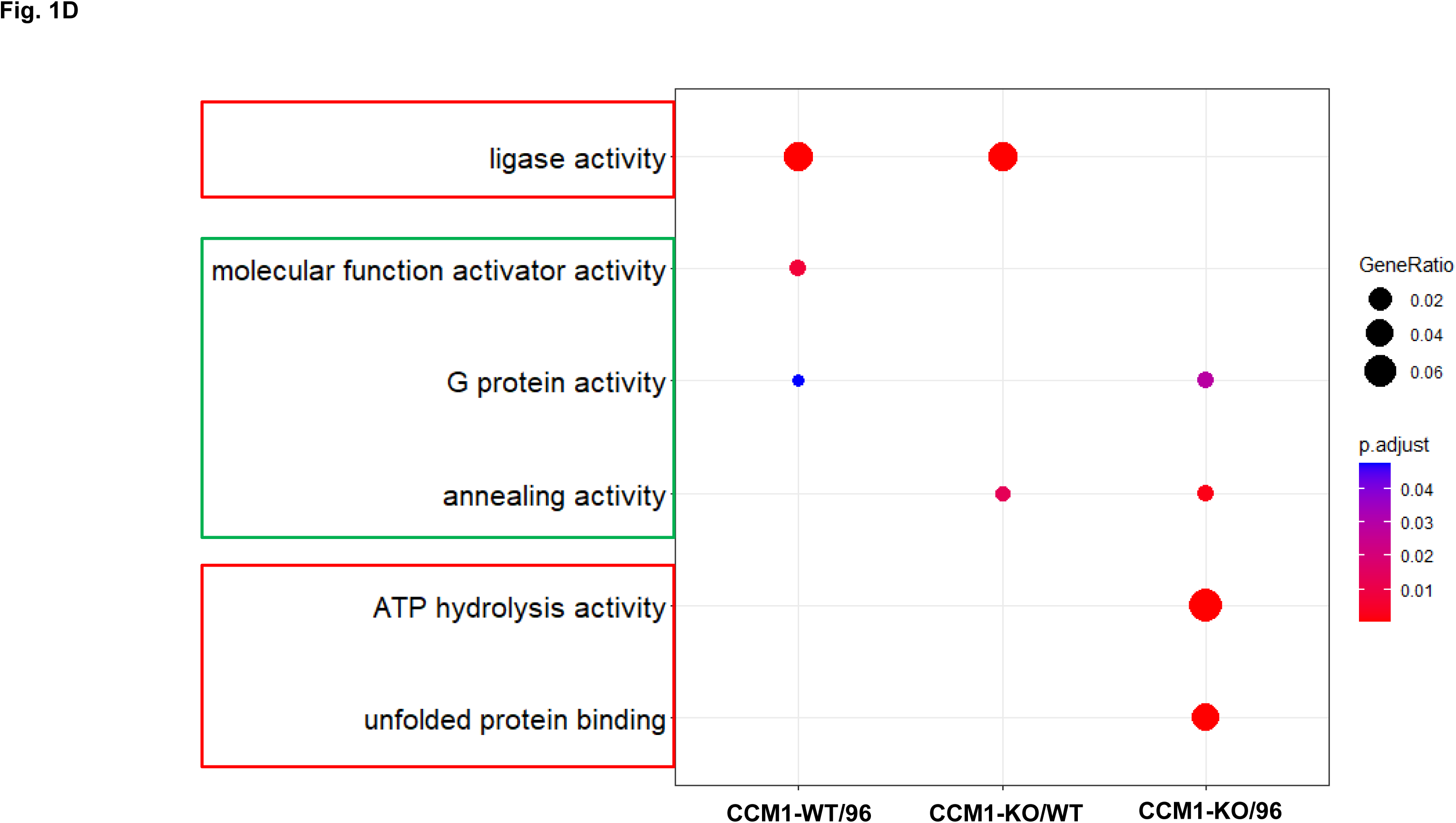
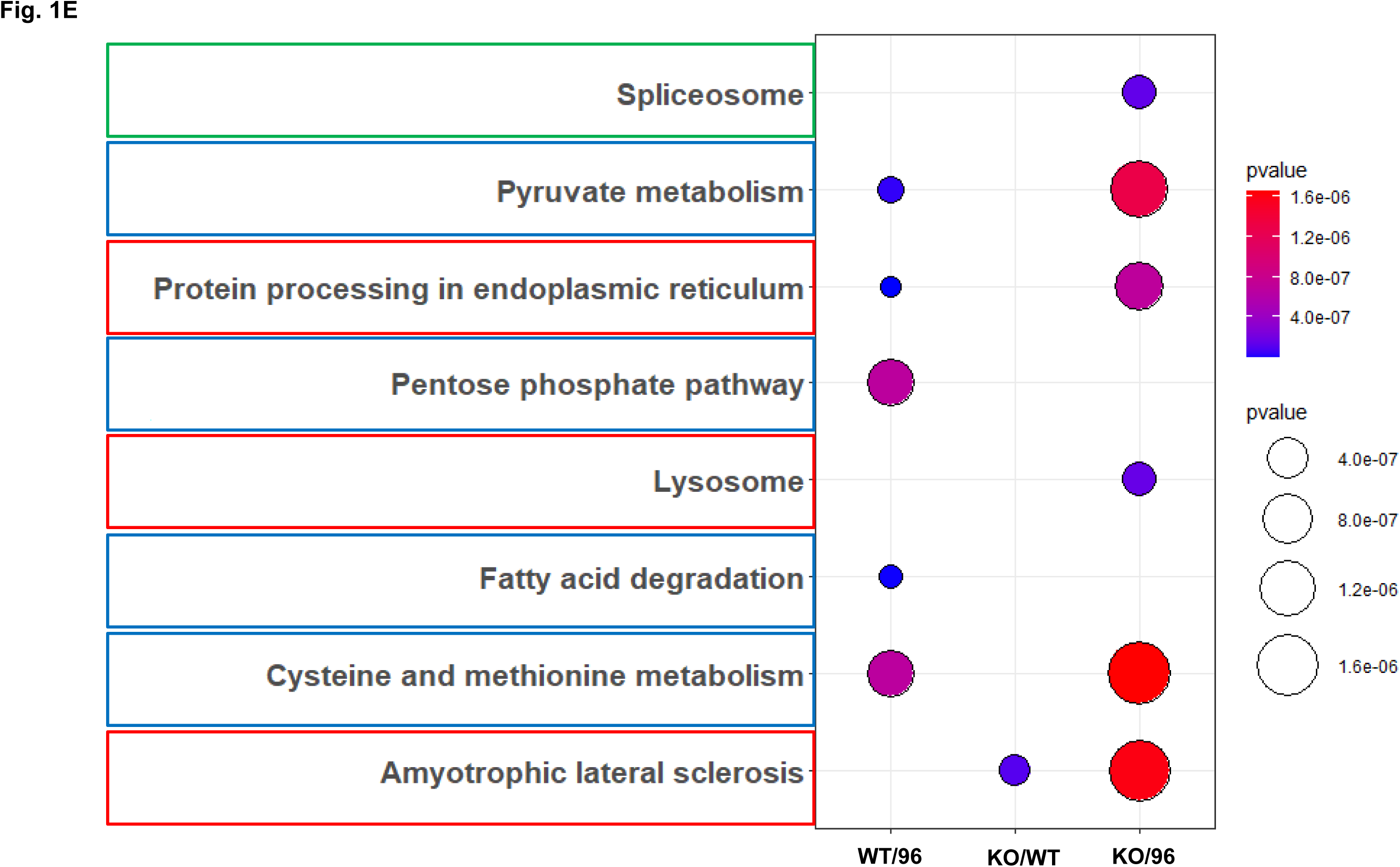
Differentially expressed protein (DEP) profiles in the MEFs with three different genotypes of CCM1, including depletion (knockout, KO), endogenously low (WT), and ectopically excessive expression (knockin, KI/96) of CCM1. The protein profiles of mouse embryonic fibroblasts (MEFs) were examined for differential expression in relation to CCM1 depletion (knockout, KO), endogenous low levels (WT), and excess expression through knockin (KI/96). These DEPs were analyzed to investigate the impact of genetic backgrounds on protein expression in MEFs. A. To compare the differentially expressed protein (DEP) profiles of three pairs of mouse embryonic fibroblasts (MEFs) with total depletion (CCM1-KO), endogenous low levels (CCM1-WT), or excess expression through knockin (CCM1-KI/96) of CCM1, a Venn diagram was utilized. This diagram illustrates the number of DEPs with significant expression changes between the paired-comparisons of CCM1 genotypes, as well as the number of specifically expressed proteins between two different CCM1 genotypes. B. To depict the transcriptome profiles of differentially expressed proteins (DEPs) in mouse embryonic fibroblasts (MEFs) across three comparative pairs of CCM1 genotypes, a bar diagram was employed. The diagram showcases the number of up-regulated (represented by red bars) and down-regulated (represented by blue bars) DEPs between two different CCM1 genotypes. C. To visualize the differential expression levels and profiles of differentially expressed proteins (DEPs) in three comparative pairs of CCM1 genotypes in mouse embryonic fibroblasts (MEFs), a Heatmap was utilized. The Heatmap displays the up-regulation (represented by red lines) and down-regulation (represented by blue lines) status of DEPs for the comparative genotype pairs across the three different CCM1 genotypes. The DEPs Heatmap was created using t-test statistical analysis and visualized with cluster software. D. The GSEA (Gene Set Enrichment Analysis) results showcasing core enriched differentially expressed proteins (DEPs) in GO pathways were presented for progesterone (PRG) treatment and vehicle control across three CCM1 genotypes. These genotypes include total depletion of CCM1 (CCM1-KO, Left), endogenous low levels of CCM1 (CCM1-WT, Middle), and ectopic overexpression of CCM1 through CCM1-knockin (CCM1-KI/96, Right) E. The GSEA (Gene Set Enrichment Analysis) results showcasing core enriched differentially expressed proteins (DEPs) in KEGG pathways were presented for progesterone (PRG) treatment and vehicle control across three CCM1 genotypes. These genotypes include total depletion of CCM1 (CCM1-KO, Left), endogenous low levels of CCM1 (CCM1-WT, Middle), and ectopic overexpression of CCM1 through CCM1-knockin (CCM1-KI/96, Right)

#### Differential signal pathways in the MEFs with different CCM1 expression levels

To uncover the biological significance behind the differentially expressed proteins (DEPs), pathway enrichment analyses were performed to group genes and establish affected pathways. However, due to the limited number of peptides identified and incomplete pathway coverage, protein data may not fully capture pathway activity compared to other high-throughput technologies, and there is a significant dataset-dependent impact on the performance of different pathway enrichment approaches (38). Therefore, the integrative use of multiple pathway enrichment analysis with different databases is highly recommended (39). In this project, both gene ontology (GO) and Kyoto Encyclopedia of Genes and Genomes (KEGG) pathway enrichment analyses were utilized. GO and KEGG pathway enrichment analyses were performed in this project. GO pathway enrichment data was further analyzed using proteomic gene set enrichment analysis (GSEA) and Upset plots (Suppl. Fig. 1A-D). Comparative GSEA plots illustrated enriched pathways among MEFs with different CCM1 protein expression levels (Fig. 1D,). Similarly, a KEGG GSEA plot provided a comparative view of enriched KEGG pathways among MEFs with varying CCM1 expression (Fig. 1E). The results revealed that protein folding, processing, and degradation pathways were the main enriched pathways influenced by cellular CCM1 protein levels. This finding was consistent in both GO and KEGG pathway analyses (red-framed pathways, Figs. 1D, 1E). Another enriched pathway involved amino acid, carbohydrate, and lipid metabolism (blue-framed pathways, Fig. 1E). Additional pathways that were identified are associated with signaling related to cellular activity (highlighted by green frames in Figs. 1D and 1E). The utilization of GO and KEGG pathways in the functional enrichment analysis yielded significant findings. These insights helped shed light on the functional differences between MEFs exhibiting different levels of CCM1 expression, allowing us to understand the underlying mechanisms and implications of these alterations (Fig. 1D, 1E). Furthermore, this analysis served as an initial filter to exclude the “CCM1 protein effect” in our subsequent omics analysis. Notably, the analysis uncovered a higher proportion of genes exhibiting down-regulation in the pathways that were identified.

### Comparative multi-omics between mouse embryonic fibroblasts with different CCM1 genotypes under progesterone (PRG) actions

This experiment aimed to analyze differentially expressed protein and RNA profiles in MEFs with distinct CCM1 genotypes in response to membrane progesterone receptor (mPR)-specific progesterone actions by comparing the progesterone (PRG) treated and control groups.

#### Differentially expressed protein (DEP) profiles in the MEFs with different CCM1 genotypes, in response to progesterone (PRG) actions

The Venn diagram analysis showed that 661 differentially expressed proteins (DEPs) were identified in the comparative pair between progesterone (PRG) treated and vehicle control groups among three different CCM1-genotypes: MEFs with endogenously low expression (CCM1-WT), total depletion (CCM1-KO), and ectopically excessive expression (CCM1-KI/96) of CCM1. Among these, 212 proteins (20%) were unique to the CCM1-KO genotype (Fig. 2A, yellow circle). Additionally, 218 DEPs (20.6%) were identified in the CCM1-WT genotype (Fig. 2A, blue circle), with 115 DEPs (10.9%) being specific to the CCM1-KI/96 genotype (Fig. 2A, pink circle). Out of the 1059 total identified proteins, 12 (1.1%) were shared by all three genotypes, 48 (4.5%) were shared by the CCM1-WT and KO genotypes, 29 (2.7%) were shared by the CCM1-KO and KI/96 genotypes, and 27 (2.5%) were shared by the CCM1-WT and KI/96 genotypes (Fig. 1A). The DEPs were further classified as up-regulated (red-color bar) and down-regulated (blue-color bar) in each CCM1 genotype under progesterone (PRG) actions. The results showed that more upregulated genes were exclusively identified in the CCM1-KO genotype, while more downregulated genes were observed in the CCM1-WT and CCM1-KI/96 genotypes (Fig. 2B). Under progesterone (PRG) treatment, a heatmap showcased substantial variations in protein expression profiles, specifically highlighting the differentially expressed proteins (DEPs) across MEFs with three distinct CCM1 genotypes. These findings underscored the significant disparities in protein expression levels among the three CCM1 genotypes (Fig.2C).

**Figure 2.**
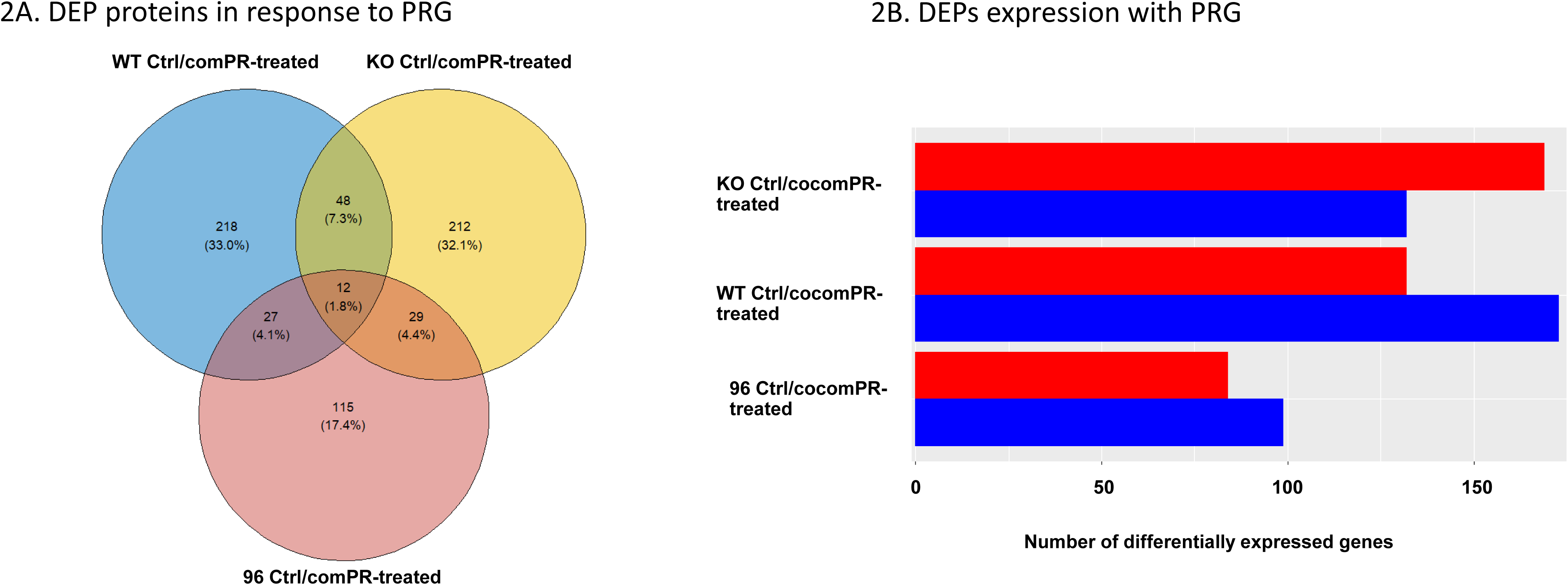
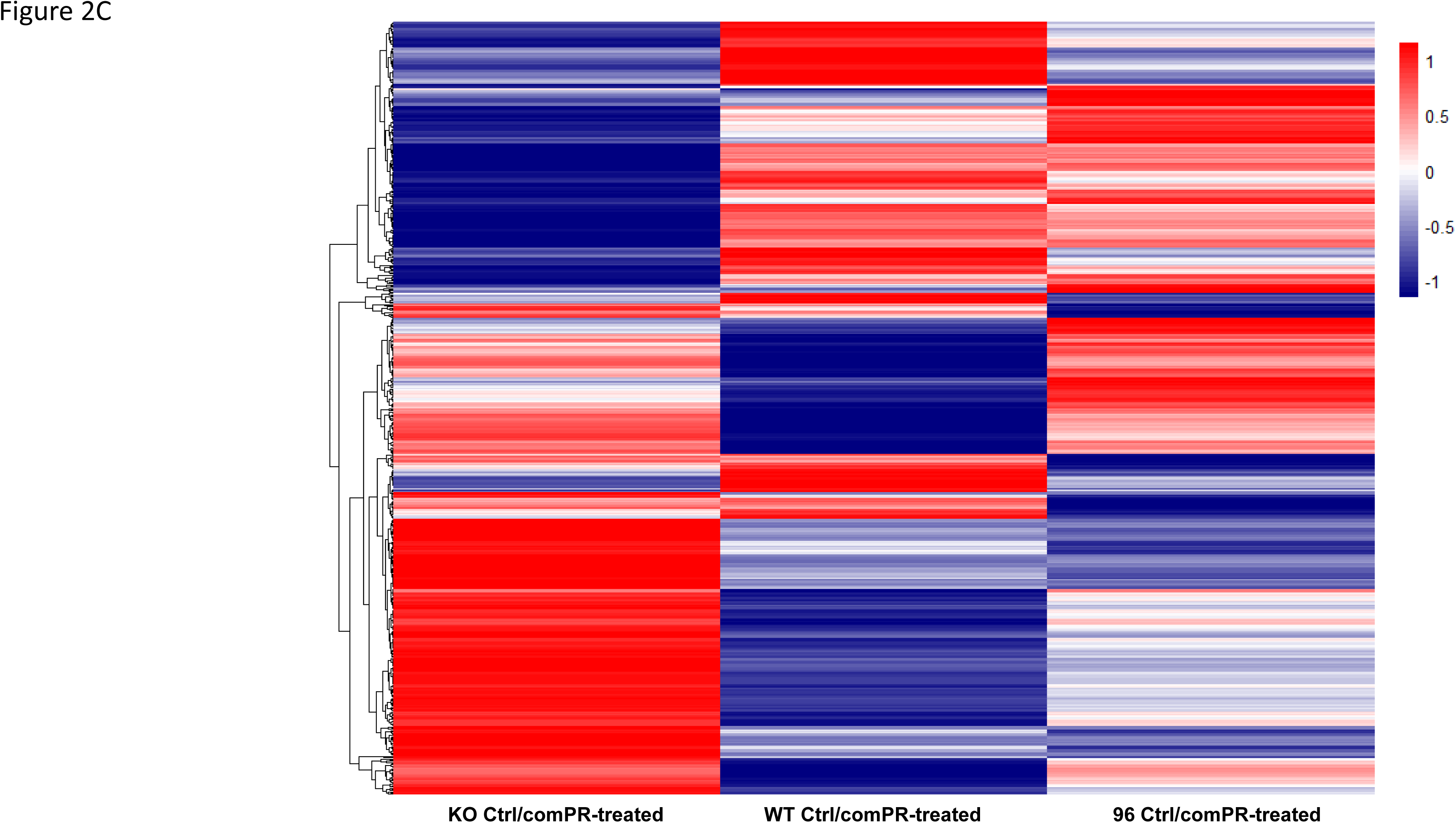
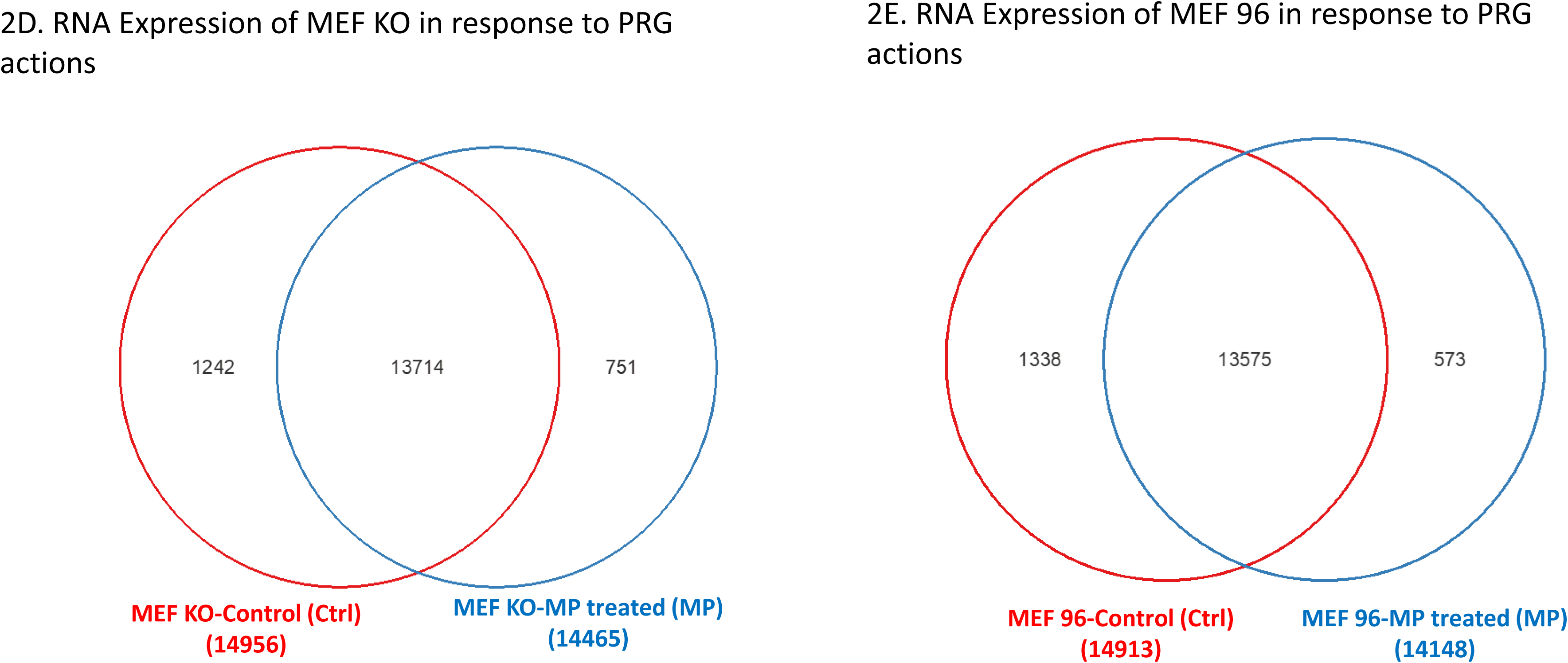
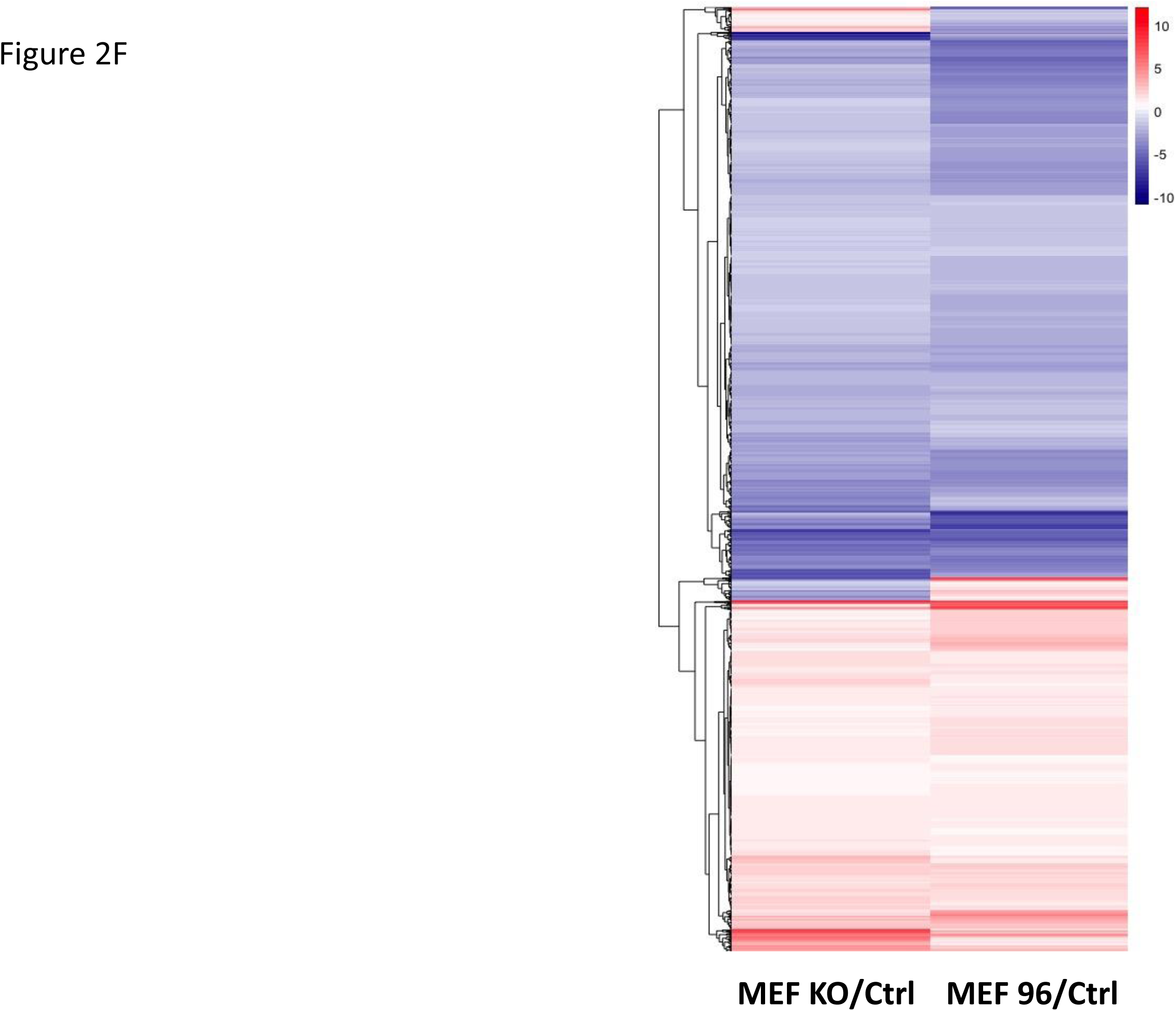
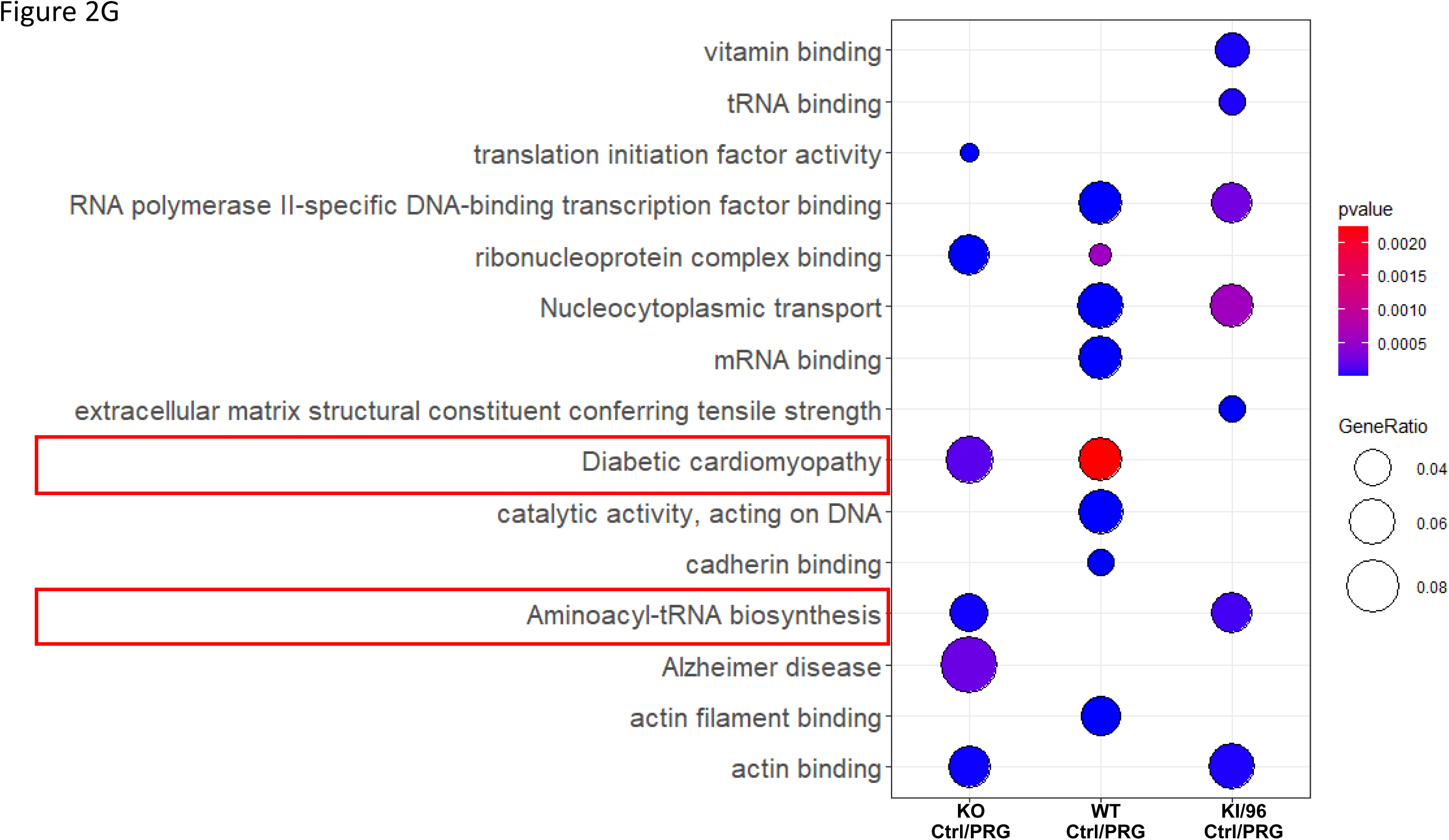
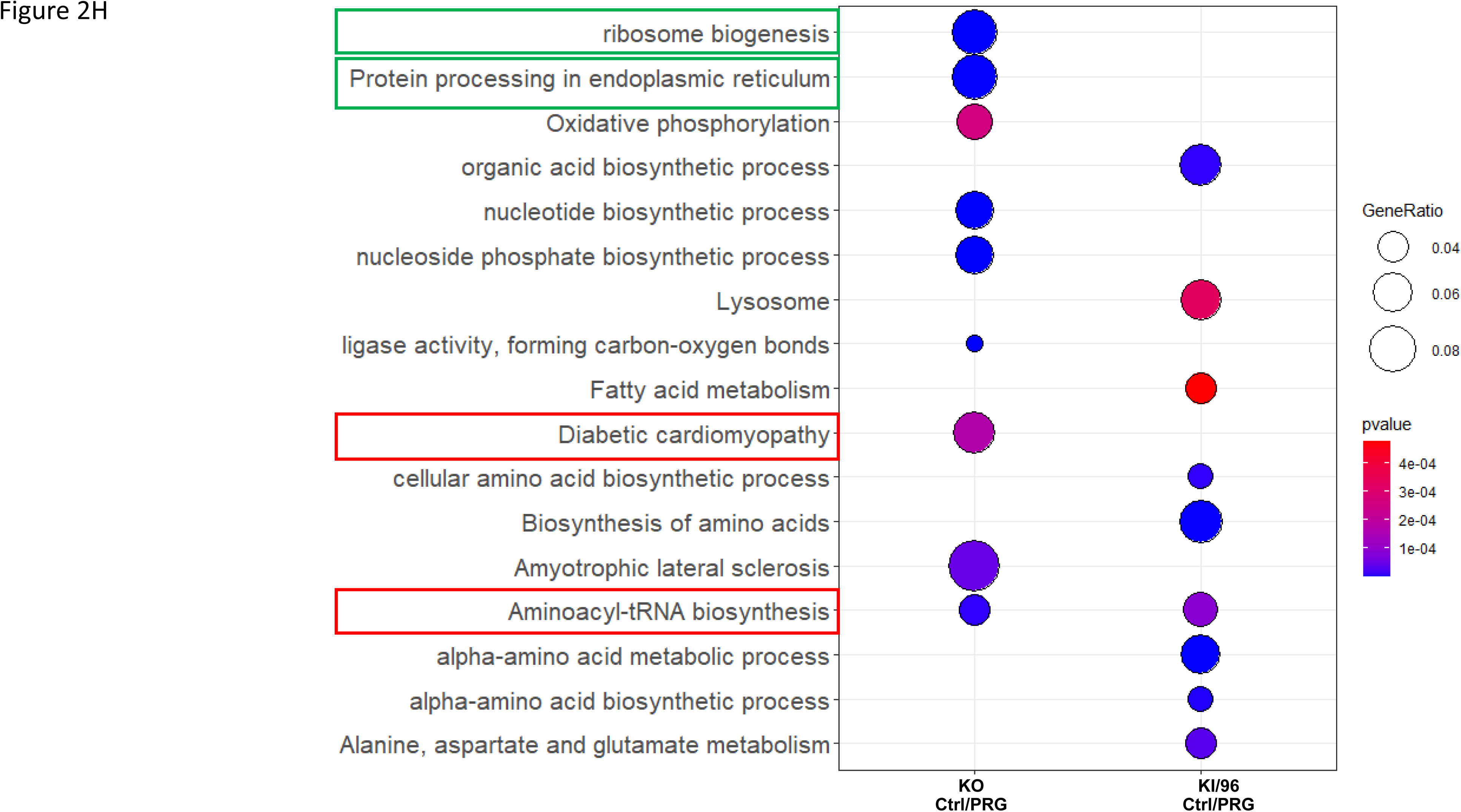
A multiomics approach to analyze Proteomic and RNA-Seq data under various levels of CCM1 expression in response to progesterone treatments. Both protein and RNA expression profiles of MEFs were investigated across different levels of CCM1 expression. The proteomic data included three distinct genotypes: complete depletion of CCM1 through CCM1-knockout (CCM1-KO), natural low levels of CCM1 (CCM1-WT), and ectopic overexpression of CCM1 through CCM1-knockin (CCM1-96). On the other hand, the RNAseq data comprised two levels of expression profiles: depleted (Knock out) and ectopic overexpression of CCM1 through CCM1-knockin (CCM1-96). **A.** The protein profiles of differentially expressed proteins (DEPs) in response to progesterone (PRG) actions were investigated in MEFs with three different CCM1 genotypes. A Venn diagram was used to display the number of specifically DEPs between progesterone (PRG) treatment and vehicle controls among the three CCM1 genotypes. The protein profiles of differentially expressed proteins (DEPs) were examined in mouse embryonic fibroblasts (MEFs) with three distinct CCM1 genotypes in response to progesterone (PRG) actions. A Venn diagram was utilized to depict the number of DEPs that were specifically affected by progesterone (PRG) treatment in comparison to vehicle controls among the three CCM1 genotypes. **B**. To illustrate the profiles of differentially expressed proteins (DEPs) in response to progesterone (PRG) actions across three CCM1 genotypes, a bar diagram was employed. The diagram provides a detailed distribution of up-regulated DEPs (represented by a red color bar) and down-regulated DEPs (represented by a blue color bar). **C.** A heatmap was used to display the profiles of differentially expressed proteins (DEPs) in response to progesterone (PRG) actions across three CCM1 genotypes. The heatmap provides a detailed distribution of up-regulated DEPs (represented by a red color line) and down-regulated DEPs (represented by a blue color line). The comparative analysis of DEPs was conducted using a t-test statistical analysis. **D.** For RNA expression profiling, Venn Diagrams were employed to compare the expression patterns between untreated MEF KO control and MEF -KO treated with progesterone (PRG) in CCM1-depletion genotype (MEF KO). The analysis revealed the presence of 751 unique differentially expressed genes (DEGs) when exposed to PRG. **E.** Likewise, in the case of ectopic overexpression of CCM1 (MEF 96) genotype, we compared the expression patterns between untreated MEF 96 control and MEF 96 treated with progesterone (PRG). Through this analysis, we discovered 573 differentially expressed genes (DEGs) when exposed to PRG. **F.** A heatmap illustrates the profiles of differentially expressed genes (DEGs) in response to progesterone actions across two CCM1 genotypes. A comprehensive analysis involved examining 1900 shared RNAseq profiles between the two CCM1 genotypes to explore how the exposure to PRG impacts the regulation of common sequences within different levels of CCM1, both treated with and without progesterone. **G.** The GSEA (Gene Set Enrichment Analysis) results showcasing core enriched differentially expressed proteins (DEPs) in GO and KEGG pathways were presented for progesterone (PRG) treatment and vehicle control across three CCM1 genotypes. These genotypes include total depletion of CCM1 (CCM1-KO, Left), endogenous low levels of CCM1 (CCM1-WT, Middle), and ectopic overexpression of CCM1 through CCM1-knockin (CCM1-KI/96, Right). The plot displayed in the analysis was specifically filtered by CCM1-associated pathways, with the original data provided in the supplementary materials. It is important to note that all samples underwent triplicate analysis, and statistical significance was determined using a student’s t-test, considering p-values less than 0.05 as significant. **H**. The RNAseq data was subjected to GSEA (Gene Set Enrichment Analysis) similar to the previous approach used for proteomic data. Notably, we highlighted the shared pathways between the RNAseq profile and the proteomic data by denoting them with bold borders, specifically focusing on Diabetic cardiomyopathy and Aminoacyl-tRNA biosynthesis.

#### Differentially expressed gene (DEG) profiles in the MEFs with two opposite CCM1 genotypes, in response to progesterone (PRG) actions

Similarly, RNAseq experiment generated differentially expressed genes (DEGs) in MEFs with two opposite CCM1 genotypes, classified as CCM1-total depletion genotype (CCM1-KO), and ectopically overexpressed CCM1 genotype (CCM1-KI/96). These genotypes exhibited significantly binary ON/OFF expression of CCM1 protein. To investigate the effects of two opposite expression pattern of CCM1 on the responsive DEG profiles in MEFs under membrane-specific (mPR-specific) progesterone (PRG) actions. In this comparative genomic analysis, a Venn diagram was firstly used to identify differentially expressed genes (DEGs) in two opposite CCM1 genotypes, classified as CCM1-total depletion genotype (CCM1-KO), and ectopically overexpressed CCM1 genotype (CCM1-KI/96). The analysis using Venn diagrams showed that when treated with membrane-specific (mPR-specific) progesterone (PRG) actions, a total of 15,707 genes exhibited differential expression in the CCM1-KO genotype pair (Fig. 2D). Conversely, in the CCM1-KI/96 genotype with ectopically excessive expression of CCM1, 15,486 genes showed differential expression (Fig. 2E). The HeatMap plot provided a visual representation of the variation in gene expression between MEFs with two different CCM1 genotypes under progesterone (PRG) treatment, indicating significant differences in gene expression profiles (Fig. 2F). These findings suggest that the presence of the CCM1 gene may enhance the induction of differentially expressed genes (DEGs) through the specific action of mPR-mediated PRG signaling.

#### Different pathway response in the MEFs with distinct CCM1 genotypes under progesterone (PRG) actions

To investigate the effects of varying levels of CCM1 on the responsive DEP profiles in MEFs under progesterone (PRG) actions. We need to understand the unique pathways involved by comparing MEFs with depleted (CCM1-KO), endogenously low (CCM1-WT), or ectopically excessive expression of CCM1 (CCM1-96) under progesterone (PRG) treatments compared to vehicle controls. To accomplish this, we employed GO pathway enrichment data and conducted proteomic gene set enrichment analysis (GSEA) (Suppl. Fig. 2C, 2D). We also performed comparative GO GSEA to examine the differentially expressed proteins (DEPs) in progesterone (PRG)-treated MEFs compared to their corresponding vehicle controls, considering different genotypes of CCM1 (various levels of CCM1 protein expression) (Fig. 2G, Suppl. Fig. 2C). Additionally, we employed a comparative KEGG GSEA plot to visualize the enriched pathways among MEFs with distinct levels of CCM1 protein expression (Fig. 2H, Suppl. Fig. 2D, 2E, 2F).

Our findings demonstrated that upon removing the initially enriched pathways linked to CCM1 expression, such as protein folding, processing, and degradation pathways (highlighted in red frames in Figs. 1D and 1E), along with the common amino acid, carbohydrate, and lipid metabolism pathway (highlighted in blue frames in Fig. 1E), there were notable enrichments observed in transcriptional and translational signal pathways in response to progesterone (PRG) actions. These findings suggest that the underlying mechanisms of CCM1 protein in the CmPn signaling network in response to progesterone (PRG) actions involve both transcriptional and translational signal pathways.

### The collection of proteomic data can be leveraged to create a pathway filter specifically for non-mPR progesterone (PRG) actions modulated by CCM1 within the CmPn network

The glucocorticoid receptor (GR) and nuclear progesterone receptor (nPR) share the same DNA sequence in their promoter regions (40), and also have a common antagonist, mifepristone (MIF or RU486), as their ligand (41, 42). This has led to the application of MIF in various clinical therapies (41–46). Numerous cell types have been found to express both the glucocorticoid receptor (GR) and nuclear progesterone receptor (nPR) (40, 47–52). The combined actions of the classic and non-classic progesterone (PRG) receptors (nPR/mPR) have been studied through the use of progesterone treatment (23, 25, 26, 52–56). There exists only one source of well-established proteomic data for U2OS cells treated with Mifepristone (RU 486) (57). U2OS cells, a human sarcoma with epithelial origin that expresses both GR and nPR (58, 59), which can undergo EMT to link with fibroblasts (60). MEFs, fibroblasts that also express both GR and nPR, are similarly affected by Mifepristone (57, 61–63). This is because Mifepristone acts as an antagonist for both GR and nPR (57–63). In this section, we will utilize the existing dataset to construct a filter aimed at excluding any “antagonist” effects caused by non-mPR-specific progesterone (PRG) actions, such as PRG actions mediated by the glucocorticoid receptor (GR) and nuclear progesterone receptor (nPR) (see Supplementary Figure 3A, 3B).

Through Venn diagram analysis, we observed that under progesterone (PRG) treatments, across MEFs with three different CCM1 genotypes (CCM1-WT with low expression, CCM1-KO with total depletion, and CCM1-KI/96 with excessive expression), a total of 661 differentially expressed proteins (DEPs) were identified after undergoing filtration (see Supplementary Figures 3A and 3B) (Supplement Figs. 3B). Among these membrane progesterone receptor (mPR)-specific DEPs, 224 (25.6%) were unique to the CCM1-KO genotype (Supplemental Fig. 3B, yellow circle), 262 (29.9%) were identified in the CCM1-WT genotype (Supplemental Fig. 3B, blue circle), and 151 (17.2%) were specific to the CCM1-KI/96 genotype (Supplemental Fig. 3B, pink circle). Out of the 876 filter-selected (mPR-specific) DEPs, 52 (5.9%) were common to all three genotypes, 79 (9.0%) were shared by the CCM1-WT and KO genotypes, 42 (4.8%) were shared by the CCM1-KO and KI/96 genotypes, and 66 (7.5%) were shared by the CCM1-WT and KI/96 genotypes (Supplemental Fig. 3B). The heatmap depicted the significant differences in DEPs among MEFs with three different CCM1 genotypes under progesterone (PRG) treatment, highlighting the distinct protein expression profiles after filter selection (Supplemental Figure 3F). The volcano plots showed the up-regulated (red dots on the right) and down-regulated (blue dots on the left) DEPs in MEFs with CCM1-KO genotype (left panel), CCM1-WT genotype (middle panel), and CCM1-KI/96 genotype under progesterone treatment (PRG) (Supplemental Fig. 3D). After passing through the filter with the CCM1 protein (red-framed pathways, Suppl. Figs. 2C, 2E), a comparative GSEA plot that integrated both GO and KEGG analyses illustrated the enriched pathways through the filtration. The identified non-mPR-specific pathways modulated by CCM1 protein, including DNA replication and DNA repair pathways (involving single-stranded DNA helicase activity, helicase activity, DNA helicase activity, and catalytic activity acting on DNA), along with cell-cell adherent junctions (cadherin binding), are prominently associated with cell proliferation (Fig. 3A, Supplementary Figures 3H-J). Likewise, a similar approach was applied to RNAseq data to generate the nuclear progesterone receptor (nPR) response filter. This filter identified GO/KEGG enriched pathways, including angiogenesis (highlighted in black), tumorigenesis (highlighted in red), cell proliferation and performance (highlighted in green), RNA process (highlighted in light blue) and inflammatory response (highlighted in purple) (Fig. 3B, Supplementary Figures 3K), serving as the non-mPR specific progesterone (PRG) actions filter for RNAseq data. These findings indicate that the majority of the identified pathways associated with PRG actions and regulated by CCM1 (Figs. 2G, 2H) fall into the second filter, suggesting their dominant involvement in non-mPR specific PRG actions within MEF cells.

**Figure 3.**
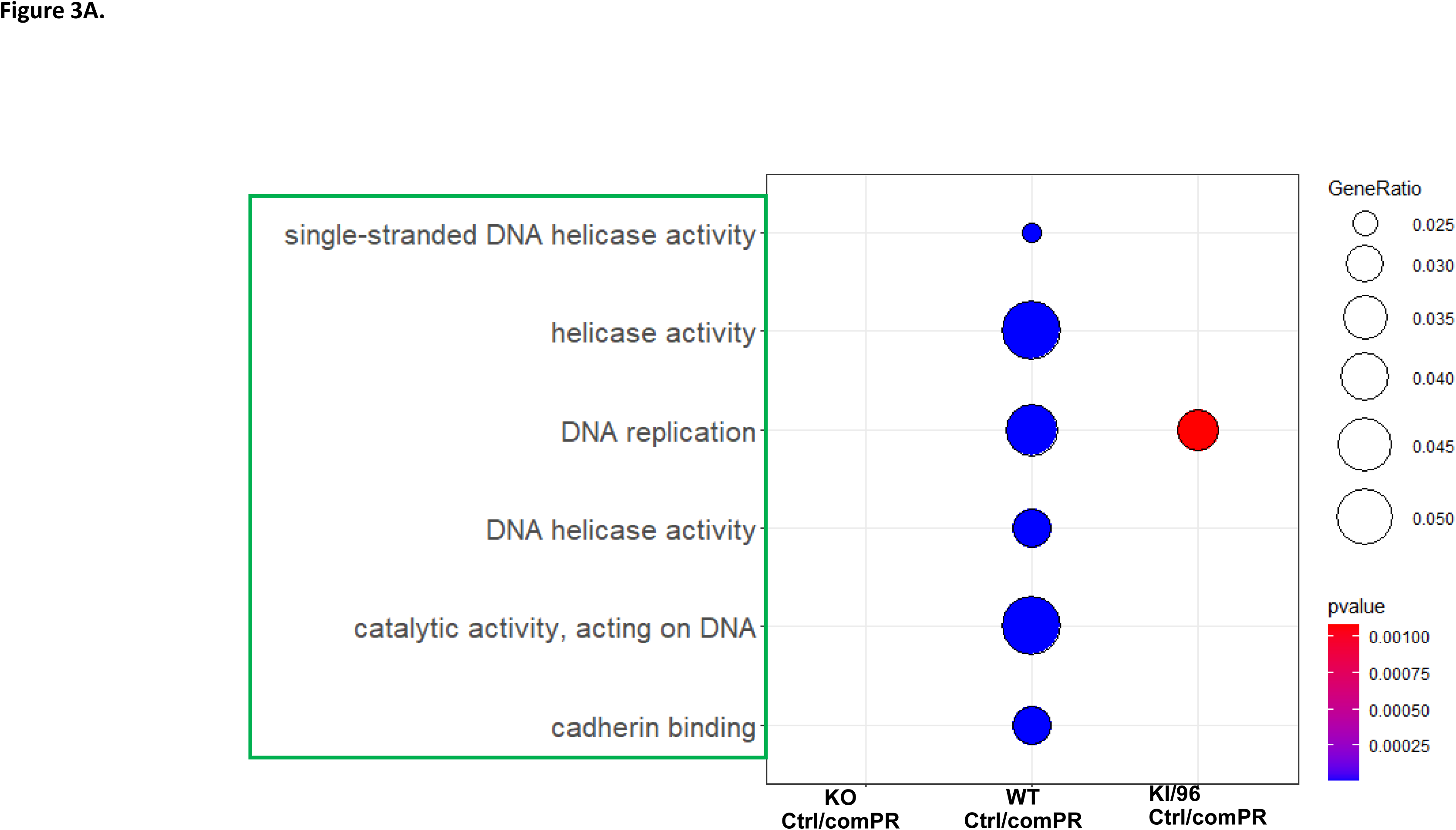
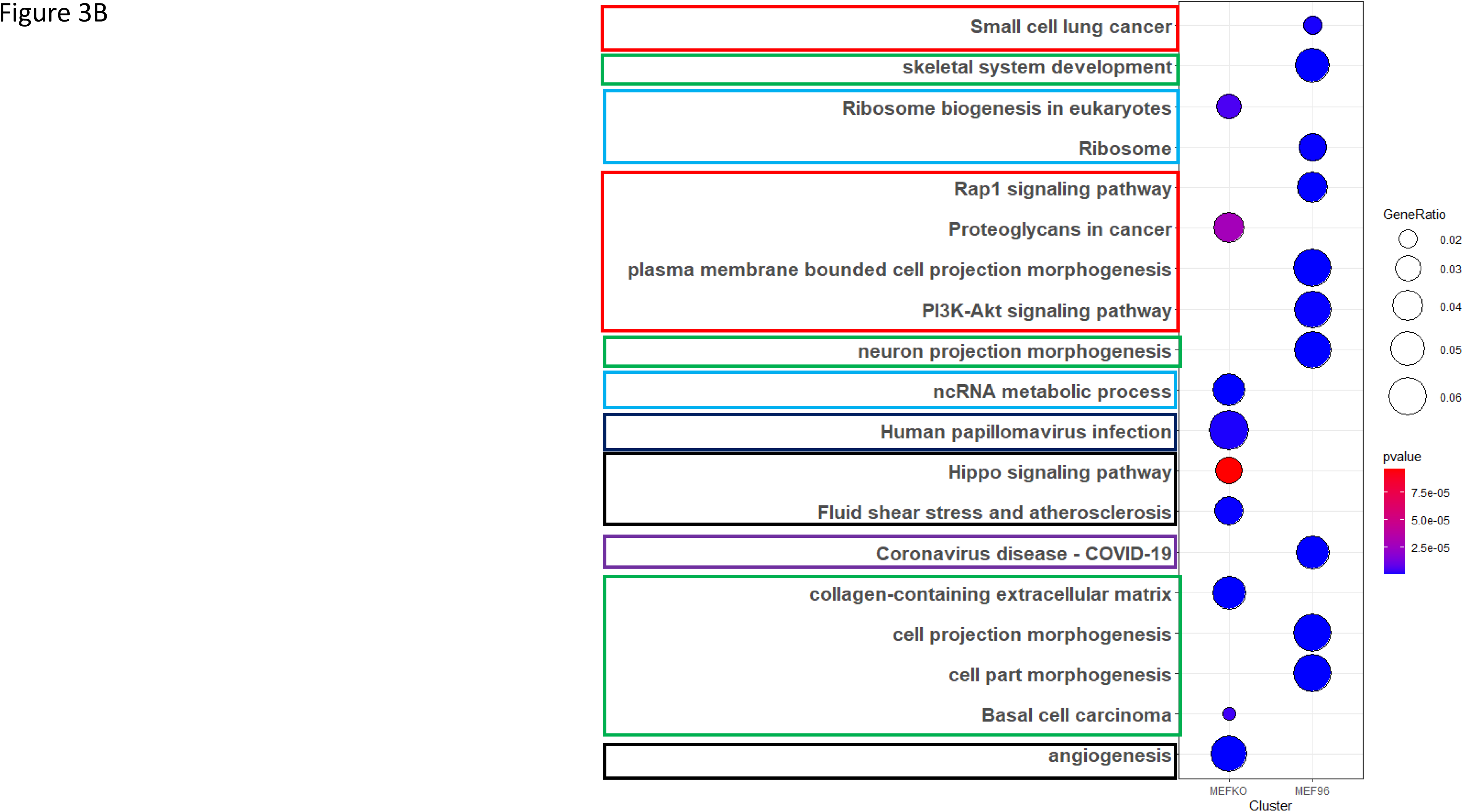
Pathways associated with progesterone (PRG) actions that are not specific to mPR among three MEF genotypes of CCM1. Considering the available data, our initial focus was on progesterone (PRG) actions that are not mediated by mPR. We accomplished this by employing the filter dataset from mifepristone (MIF, as an antagonist) to identify pathways shared by the glucocorticoid receptor (GR) and the classic nuclear progesterone receptor (nPR). It’s important to note that MIF can only function as an agonist and works synergistically with PRG on mPR-mediated signaling. A. The GSEA (Gene Set Enrichment Analysis) results showcasing core enriched differentially expressed proteins (DEPs) in GO/KEGG pathways were presented for progesterone (PRG) treatment and vehicle control across three CCM1 genotypes. These genotypes include total depletion of CCM1 (CCM1-KO, Left), endogenous low levels of CCM1 (CCM1-WT, Middle), and ectopic overexpression of CCM1 through CCM1-knockin (CCM1-KI/96, Right) **B.** The GSEA (Gene Set Enrichment Analysis) results showcasing core enriched differentially expressed genes (DEGs) in GO & KEGG pathways were presented for progesterone (PRG) treatment and vehicle control across two CCM1 genotypes. These genotypes include total depletion of CCM1 (CCM1-KO, Left) and ectopic overexpression of CCM1 through CCM1-knockin (CCM1-KI/96, Right)

### Identification of mPR-specific PRG pathways modulated through CCM1 proteins in the CmPn signal network

In the previous experiments, we established the nPR-specific progesterone action filter. Through a two-step filtration process, we identified mPR-specific progesterone (PRG) pathways. The first filtration involved the pathway filter based on CCM1 protein expression levels, as defined earlier. The second filtration utilized the pathways filter for non-mPR specific PRG actions, as illustrated (Suppl. Fig. 3A). According to our findings, the actions of mPR-specific PRGs, mediated by CCM1 within the CmPn network, are less common than PRG actions through non-mPR specific PRGs, particularly the glucocorticoid receptor (GR) and nuclear progesterone receptor (nPR) (Fig. 3A, 3B). The mPR-specific progesterone (PRG) action exhibits unique sensitivity to CCM1 regulation. By analyzing proteomic data, we conducted a comprehensive examination of both GO and KEGG pathways, which revealed that the PRG pathways specific to mPR (membrane progesterone receptor) are enriched with molecular chaperones, specifically HSPs-70 and 90, that interact with steroid receptors in the absence of ligands. Moreover. Other identified pathways include axonal guidance transmembrane signaling receptors (specifically semaphorin receptor), cell-cell junctions and communications (specifically gap junction), as well as the membrane “chaperone” phospholipid (phosphatidylethanolamine, PE), which is known to contribute to multiple common signaling pathways like MAP kinase (MAPK) (illustrated in Fig. 4A). Notably, certain pathways, such as cellular autophagy (TORC2 complex) and translational activity (ribosome), were excluded during the filtering process (highlighted in a yellow frame), as indicated in the two filters (Figs. 1D, 1E, 3A, 3B). Likewise, the analysis of GO and KEGG pathways using RNAseq data revealed the enrichment of multiple pathways that are specific to the membrane progesterone receptor (mPR). Following the exclusion of false-positive pathways, such as RNA process protein folding (highlighted in a yellow frame), a prominent pathway that emerged was the calcium channel pathway (highlighted in a green frame). Additionally, other pathways identified included cellular performance (highlighted in a light blue frame) and protein synthesis (highlighted in a red frame) (shown in Fig. 4B). In our combined analysis, we conducted GSEA using GO and KEGG pathway analysis, integrating omics data. After eliminating the components from two filters (indicated by the yellow frame), we observed that the majority of pathways align with our previously described pathways (highlighted in green), including the prominent calcium channel pathway, molecular chaperones, axonal guidance pathway, and cellular autophagy. However, we also identified unexpected pathways, such as inflammatory pathways (highlighted in blue), and notably, transcriptional factors (highlighted in red), suggesting the potential involvement of transcriptional regulation in CCM1-mediated mPR-specific progesterone actions (Fig. 4C).

**Figure 4.**
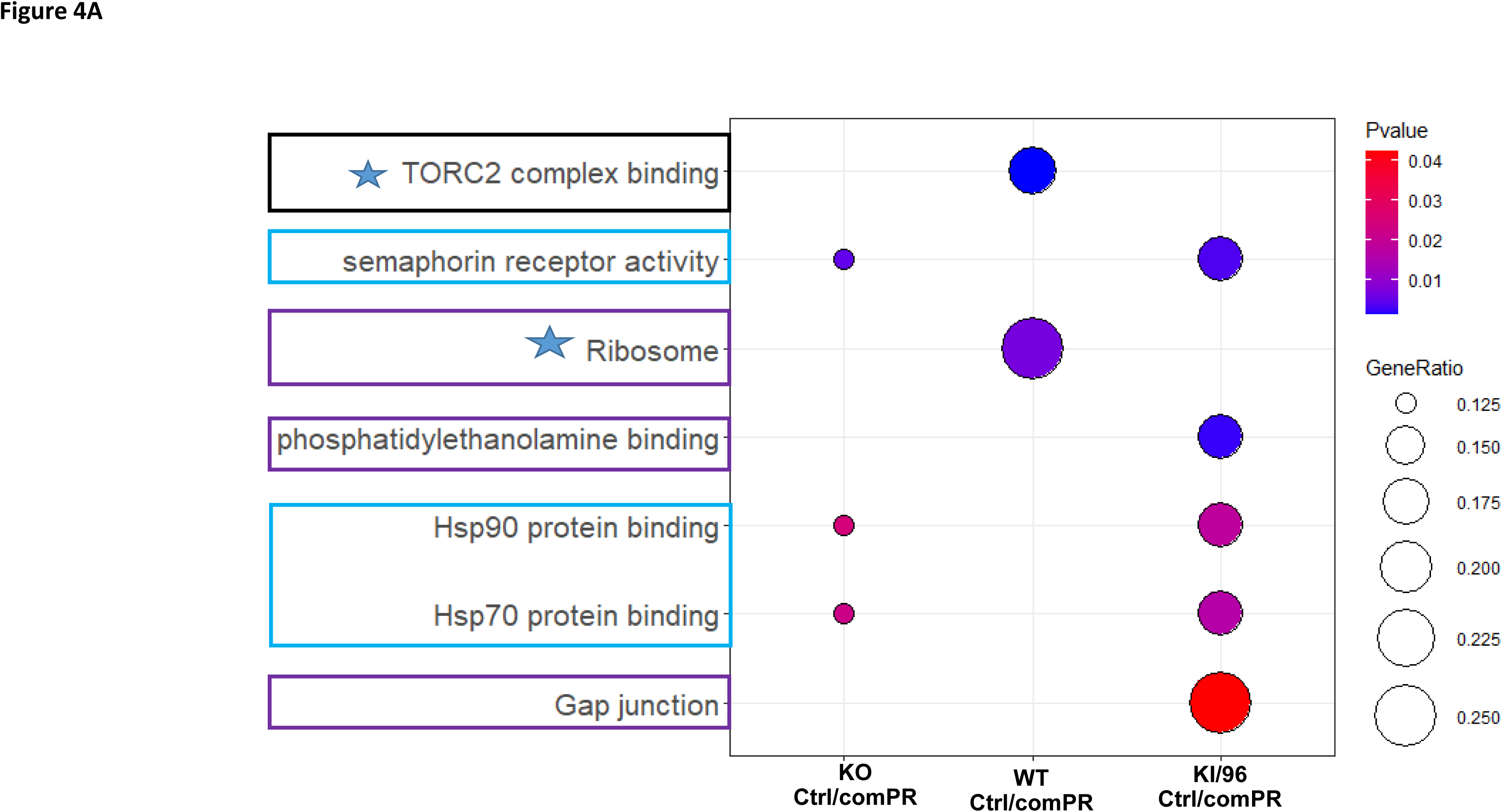
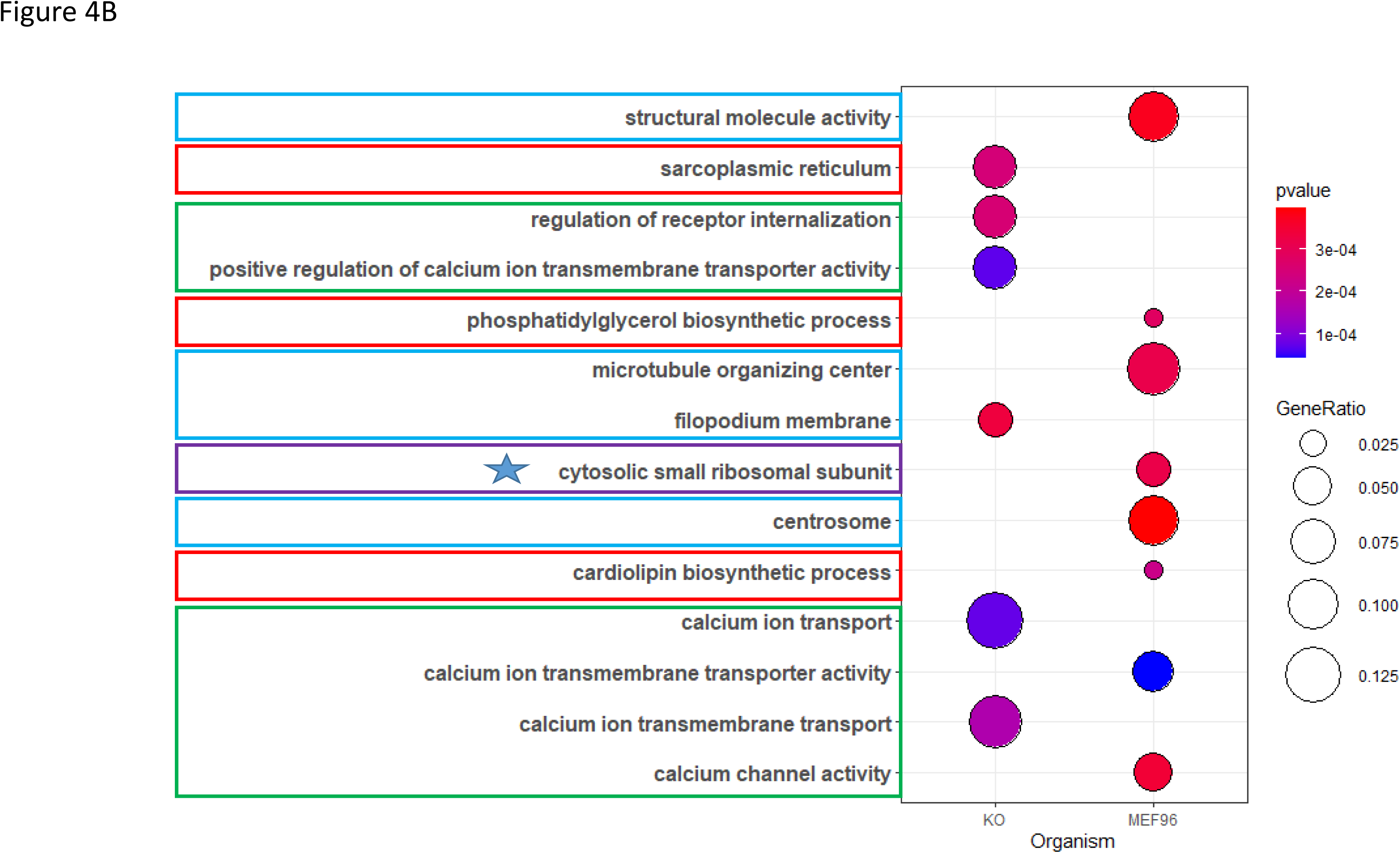
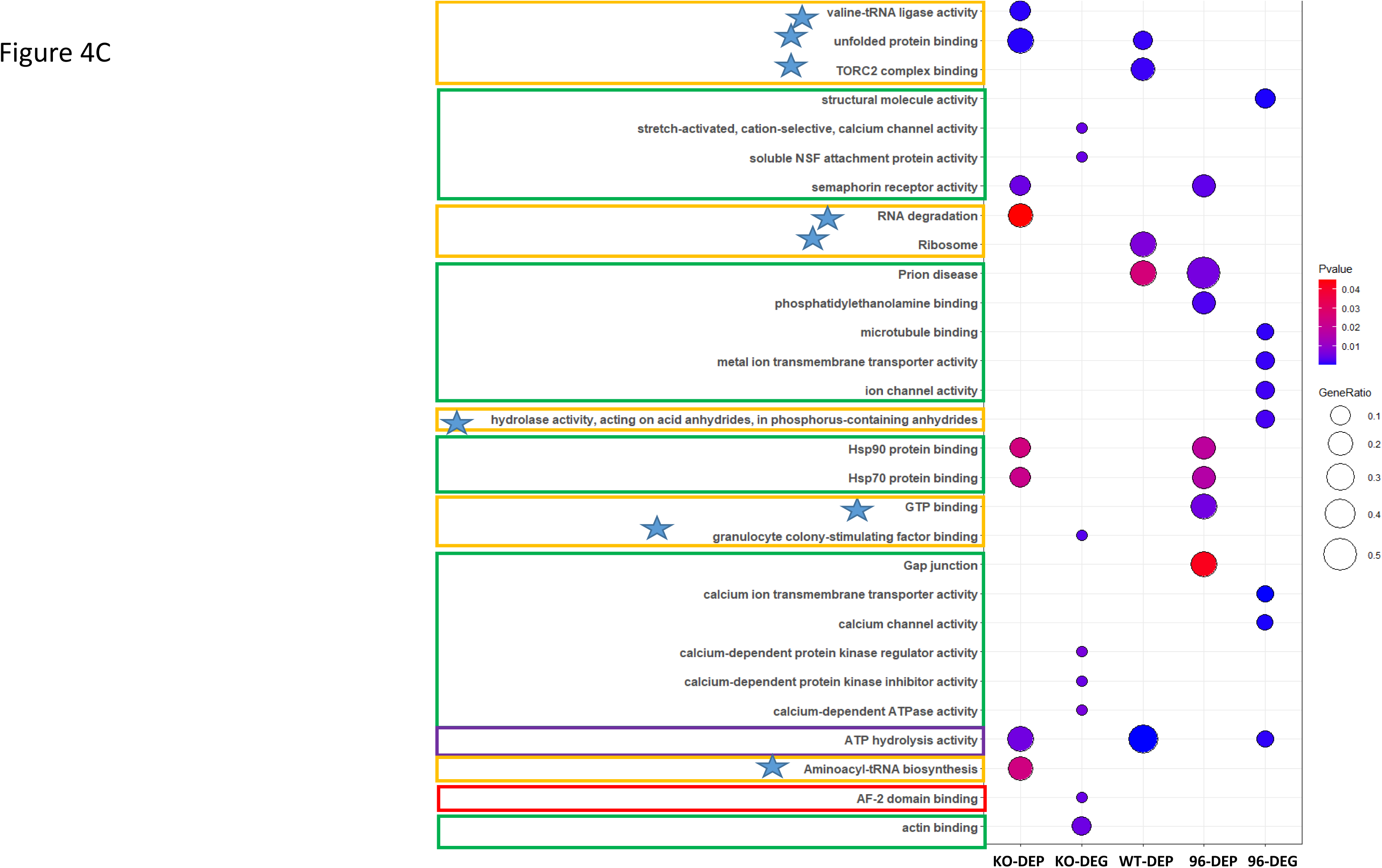
Signal pathways affected by membrane-specific (mPR-specific) progesterone (PRG) actions among MEFs with different genotypes of CCM1. The objective of this experiment was to identify signal pathways specifically associated with the membrane progesterone receptor (mPR) by excluding progesterone (PRG) actions that are not specific to mPR across the distinct CCM1 genotypes defined in the previous step. To achieve this, we utilized a filtered dataset from mifepristone (MIF), a commonly used antagonist of the classic nuclear progesterone receptor (nPR), to identify pathways that are shared by the glucocorticoid receptor (GR) and nPR, as generated in the last section. It is important to note that MIF functions solely as an agonist and exhibits synergistic interactions with progesterone (PRG) actions on mPR-mediated signaling, as described in the previous figure. **A.** To visualize the GSEA results of core enriched DEPs from GO and KEGG pathway enrichments, an integrative dot plot was used. The plot displayed the data from three CCM1 genotypes that passed the filter for non-mPR-specific PRG actions. Triplicate analysis was conducted, and statistical significance was determined using a student’s t-test with a p-value cut-off of less than 0.05. The integrative dot plot highlighted the enriched pathways associated with mPR-specific PRG action, indicated by red-framed, blue-framed, and black pathways. **B.** Similarly, an integrative dot plot was employed to visualize the GSEA results of enriched RNAseq data from GO and KEGG pathway enrichments. This data, which underwent triplicate analysis, passed through the filter for non-mPR specific PRG actions among the two CCM1 genotypes utilized in the RNAseq portion of the study. The statistical significance of the results was determined using a student’s t-test, with a p-value cut-off of less than 0.05. **C.** Finally, a summarized dot plot was generated to visualize the combined GSEA results of core enriched DEPs and DEGs from both proteomic and RNAseq data.

### Identification of mPR-specific action-induced and *CCM1-associated transcription factor (TF) using our newly developed ESM-based, Cost-Sensitive SVM (ESM-CS-SVM) Approach*

Following the significant findings from the previous experiment regarding the potential transcriptional regulation associated with the CCM1 protein-mediated membrane progesterone receptor (mPR)-specific progesterone action, our subsequent step involved employing machine-learning techniques to predict the involvement of transcription factors (TFs) in this signaling pathway. Specifically, we developed an ESM-based Cost-Sensitive Support Vector Machine (ESM-CS-SVM) model. Through the utilization of a 10-fold cross-validation approach, our machine-learning-based TF prediction model achieved impressive performance metrics, with an average F1 score of 0.9478, specificity of 0.9627, sensitivity of 0.9513, and balanced accuracy of 0.9570. These parameters demonstrate the effectiveness of our ESM-based, cost-sensitive SVM approach for predicting transcription factors from protein sequences. Our future endeavors will focus on optimizing the model and expanding its capabilities to handle diverse and complex datasets. In sum, by implementing machine learning techniques with a confidence interval exceeding 95% for specificity and sensitivity (Table 1A, 1B). From our optimized ***ESM-CS-SVM*** model, we identified multiple candidate TFs with approximately 50% of the testing epochs, such as the B cell receptor associated protein 31 (BCAP31) from proteomic data, which fall well in the newly identified inflammatory response pathway (Fig. 4C). In our final analysis for these identified TFs, we conducted GSEA using GO and KEGG pathway analysis, integrating omics data, we observed that the majority of pathways are indeed transcriptional regulatory pathways modulating our previously described pathways (highlighted in red), some TFs also control other cellular pathway, such as membrane transporter (highlighted in green), translational regulation (highlighted in blue), and energy supply (highlighted in purple),

**Table 1:** Machine learning models predicted the functionality of mPR-specific proteins and RNA sequences. **A.** The Evolutionary Scale Modeling and Support Vector Machine (ESM-SVM) models demonstrated the highest accuracy among the models employed in this study. These models were trained on a dataset consisting of 4331 unique FASTA sequences. The selection of the model was based on achieving an average accuracy of approximately 95% across all tests. **B.** In addition, Convolutional Neural Networks (CNN) were evaluated to determine if they could achieve a comparable accuracy of 95%. However, these attempts were unsuccessful. Nonetheless, we compared the results of both models to identify the sequences that were most likely associated with suggested transcriptional factor functionality.

| Testing Averages |  |
| --- | --- |
| F1 score | 0.948 |
| Specificity | 0.963 |
| Sensitivity | 0.958 |
| Accuracy | 0.961 |

B. Convolution Neural Network (CNN)
| Testing Averages |  |
| --- | --- |
| F1 Score | 0.934 |
| Specificity | 0.954 |
| Sensitivity | 0.947 |
| Accuracy | 0.950 |

## Discussion

Initially, isogenic mouse embryonic fibroblasts (MEFs) were established from E8.5 mouse embryos of WT and Ccm1-knockout (KO) mice. Additionally, Ccm1-knockin (KI) MEFs (CCM1-KI/96) were created by infecting Ccm1-KO MEFs with a lentiviral vector encoding human KRIT1 (CCM1-transduced, Lv-KRIT1) (4). By utilizing these three MEFs with varying Ccm1 gene dosages (WT, Ccm1-KO, Ccm1-9/6), it was found that CCM1 loss-of-function (LOF) leads to increased susceptibility to oxidative DNA damage, induction of DNA damage sensors and repair genes, and apoptotic response. These findings suggest that CCM1 may play a role in maintaining intracellular reactive oxygen species (ROS) homeostasis to prevent ROS-induced cellular dysfunctions (4). Moreover, CCM1 prevents upregulation of c-Jun induced by oxidative stimuli (5) and inhibits abnormal activation of the Nrf2 stress defense system and its downstream effectors HO-1 and Glo1 (7). The MEF toolset was also used to validate the discovery that CCM1 LOF is associated with increased expression of Vegfa and subsequently elevated activation of its receptor, VEGFR2. This leads to loss of barrier function by disrupting the β-catenin-VE-cadherin interaction in vascular endothelial cells (ECs) (8).

The blood-brain barrier (BBB) tightly controls molecular exchanges between the blood and the central nervous system (CNS), making it a crucial interface to understand for addressing neurological conditions, especially hemorrhagic stroke (64, 65). Despite the lack of readily available clinical agents or measures to prevent BBB leakage or repair it, numerous epigenetic mechanisms or regulators that are either protective or disruptive for BBB components have been identified, indicating potential therapeutic opportunities to meet this challenge in the future (66). Inflammatory events are commonly recognized as a significant insult to the BBB (12, 13), and steroids are often therapeutically used for inflammatory diseases and as a common agent against edema (11) in numerous human conditions (14, 15), including many BBB disorders. However, there are still controversies surrounding their use (67).

In the absence of hormone, steroid receptors as hormone-inducible transcription factors are bound to chaperones, which was identified as one of the mPR-specific pathways mediated by CCM1 within the CmPn signal network in this study (Fig. 4C). According to recent findings, a novel signaling network called the CSC-mPR-PRG-nPR/CSC-mPR-PRG (CmPn/CmP) operates within endothelial cells (ECs). The CmPn signaling network is only present in nPR(+) cells while the CmP signaling network can exist in both nPR(-) and nPR(+) cells (23–26, 52–56, 68, 69). Upon binding to progesterone, the receptor undergoes a conformational change and translocate to the nucleus as a transcriptional factor (42). Many steroid receptors share common ligands and can coordinate their physiological function by competing for the same ligand and DNA motifs as a transcriptional co-factor (50), such as mifepristone, a common antagonist for both glucocorticoid receptor (GR) and nuclear progesterone receptor (nPR).

Previous data of signal transduction modulated by the CCM signal complex (CSC), suggested that perturbed CSC after depletion of one of three CCM (CCM 1, 2, 3) genes, leads to blood vessel cell junction organization disruption (24, 26, 70, 71). Comparative omics data across multiple models also provided further evidences that the CSC can couple both classic nuclear progesterone receptor (nPR) and non-classic membrane progesterone receptor (mPR) to form a large CmPn/CmP signaling networks for progesterone mediated cellular actions (23–25, 29, 52, 54–56, 59, 69, 72), which may affect cell junction organization and lead to compromised BBB (26). In this study, we utilized three MEFs with differential expression levels of CCM1 toolset to perform proteomic analysis to elucidate Ccm1-mediated signaling, specifically through non-classic membrane progesterone receptor (mPR) within the CmPn signaling network.

Transcription factor (TF) prediction plays a vital role in gene expression analysis as it enables us to understand the regulatory mechanisms governing gene expression. TFs are proteins that bind to specific DNA sequences and control the transcription of target genes, thereby influencing gene expression levels. Predicting TFs involves computational methods that utilize sequence information, DNA binding motifs, and other relevant features to identify potential TFs within a gene set or genomic regions. By predicting TFs, we gain insights into regulatory networks, unravel the complexities of gene expression, and comprehend the functional roles of TFs in cellular processes and disease. Through the integration of modern statistical methods and artificial intelligence, specifically deep learning, we have identified BCAP31 as a promising transcription factor. BCAP31, the most abundant protein in the endoplasmic reticulum (ER) (73), is regulated by CCM genes but negatively influenced by PRG-mediated mPR actions. This discovery sheds light on previously unexplored implications in various disease contexts.

This experiment explored protein and gene expression variations in mouse embryonic fibroblasts (MEFs) with different CCM1 genotypes and their response to progesterone (PRG). Distinct protein expression profiles were observed among CCM1 depletion (knockout, KO), CCM1-endogenously low (WT), and ectopically excessive expression CCM1 (CCM1-KI/96) MEFs, with some overlapping proteins. These proteins were categorized as up-regulated or down-regulated in each genotype pair. Pathway analysis using gene ontology (GO) and Kyoto Encyclopedia of Genes and Genomes (KEGG) revealed the impact of CCM1 protein expression on protein folding, processing, degradation, metabolism, signaling, and transcriptional and translational pathways. The study also investigated gene expression differences between CCM1-KO and CCM1-KI/96 MEFs under progesterone (PRG) influence, highlighting the role of CCM1 in transcriptional and translational signaling pathways. Our findings suggest that CCM1 is involved in the signaling network associated with progesterone (PRG) actions with CmPn signal network. Furthermore, one novelty of the study was to utilize Omics data to create two layers of pathway filter for CCM1molecular function and progesterone (PRG) actions mediated by glucocorticoid receptor (GR) and nuclear progesterone receptor (nPR), which are not specific to the membrane progesterone receptor (mPR) ().

## Conclusion

The findings of this paper reveal the role of CCM1 in regulating cellular processes under different progesterone actions. The study found significant differences in gene and protein expression between different genotypes of CCM1 under progesterone actions, indicating that CCM1 plays a critical role in regulating CmPn network. The enriched pathways determined by cellular CCM1 protein dosage were mainly related to protein folding, processing, and degradation. However, after removing the CCM1 protein dosage-dependent pathways, the study identified significant enriched transcriptional and translational signal pathways in response to progesterone (PRG) actions. Next, using established filter, the study defined the unique pathways involved in mPR-specific PRG actions mediated by CCM1 protein within CmPn signal network. Finally, utilizing our own Machine-learning algorism, we generated a list of identified transcription factors (TFs) from our GSEA results, and elucidated the possible transcriptional regulatory mechanism for

**Figure 5:**
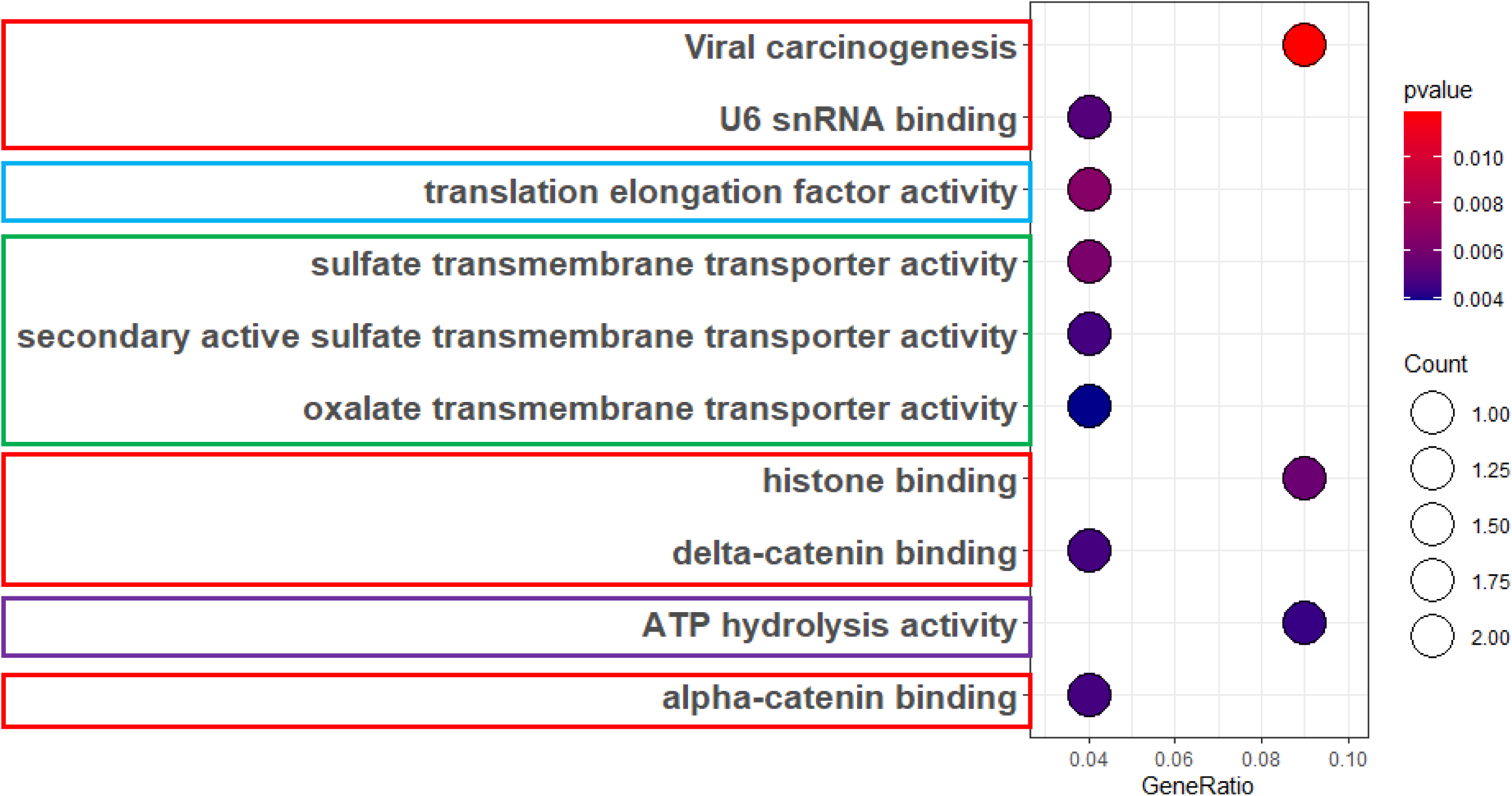
Using machine learning techniques, we identified transcriptional factors (TFs) and established novel mPR-specific regulatory pathways through the enrichment of mPR-specific DEPs and DEGs. Upon completing the machine learning-based prediction of transcription factors, we successfully identified a total of 12 potential transcription factors (TFs). To further investigate their functional roles, we utilized the Entrez ID identifiers associated with these transcription factors. By conducting Gene Ontology (GO) and Kyoto Encyclopedia of Genes and Genomes (KEGG) pathway enrichment analyses, we identified several pathways and processes in which these transcription factors potentially modulate. These findings provide valuable insights into the potential functional mechanisms and regulatory networks involving the transcriptional regulations.

## Data availability

Readers can access the data supporting the conclusions of the study through supplemental materials and some omics data are in process to deposit into NIH genomic or proteomic databases repertoire, and can be acquired by contact the corresponding author.

## Acknowledgments

We wish to thank Odalys Quintanar, Muaz Bhalli, Alexander Le, Ofek Belkin, Mellisa Renteria, David Jang, Justin Aickareth, Victoria Reid, Majd Hawwar, Revathi Gnanasekaran, Nickolas Sanchez, Charlie Harvey, and Drexell Vincent at Texas Tech University Health Science Center El Paso (TTUHSCEP); Mingchen Li and Shubham Kuppili at Texas Tech University (TTU), and Moqsadur Rahman, Cameron C Ellis, Estevao Igor, Igor C. Almeida, and Mahmud Shahriar Hossain at University of Texas El Paso (UTEP) for their technical help during the experiments.

## Author contributions

JC: Investigation, visualization, bioinformatics software analysis, data curation, writing-original draft preparation; JAF: Investigation, data validation, BG: Visualization, software, data curation, validation; LAA: Bioinformatics software analysis, data curation, LYC: Investigation, visualization, bioinformatics software analysis, data curation, VS: Conceptualization, Methodology, Investigation, reviewing, editing, JZ: Conceptualization, Methodology, Investigation, writing-original draft preparation, reviewing, editing and finalization, project supervision.

